# Structure of NO16, a marine non-tailed vibriophage with a global distribution

**DOI:** 10.1101/2025.11.28.690173

**Authors:** S. Otaegi-Ugartemendia, G. N. Condezo, M. Martínez, P. G. Kalatzis, M. Middelboe, C. San Martín

## Abstract

Non-tailed phages remain an underexplored group in marine environments, as tailed phages have dominated sequence and culture collections. However, recent surveys indicate that non-tailed phages might be more abundant than tailed phages, and with different effects on microbial mortality and gene transfer. Here, we report the structural characterization of bacteriophage NO16, a non-tailed vibriophage, one of the simplest members of the Double Jelly Roll (DJR) lineage. Mass spectrometry analyses indicate that the virion is composed of at least nine different proteins. NO16 has a *pseudo*T = 21 capsid, similar to other phages in the lineage but differs in the organization of minor capsid proteins, mainly in terms of membrane-capsid contacts. The DJR major capsid protein GP19 is stabilized by a cation in the base as previously observed in corticovirus PM2, and by strong electrostatic interactions between monomers. Notably, localized reconstruction reveals a symmetry mismatch at the vertex, where two trimeric GP13 spikes are anchored to a pentamer of the penton base GP14. GP13 presents carbohydrate-binding modules and putative glycosylase activity, indicating implications for host entry. Finally, we use structural and functional predictions for NO16 non-structural proteins to propose a complete atlas of the NO16 infectious cycle.

## Introduction

While tailed bacteriophages have traditionally been considered the most abundant viruses in the marine environment, recent surveys on marine samples suggest that non-tailed phages may actually exceed tailed phages on abundance and diversity (Brum, Schenck et al., 2013, Kauffman, Hussain et al., 2018, Steensen, Seneca et al., 2024, Yutin, Backstrom et al., 2018). Most non-tailed phages are members of the *Varidnaviria* realm (formerly Double Jelly Roll lineage). These DNA viruses encode major capsid proteins (MCP) folding as a double jelly roll (formed by eight β-sheets in each sandwich) perpendicular to the capsid surface (Koonin, Dolja et al., 2020, Krupovic, Makarova et al., 2022, Yutin et al., 2018). DJR lineage members include DNA viruses infecting hosts in the three domains of life: bacteria, archaea and eukarya.

Overall, few bacteria-infecting viruses from the DJR lineage have been structurally studied compared to tailed bacteriophages. Non-tailed bacteriophages are not easy to grow and purify in the laboratory, at least with standardized phage purification protocols (Bardy, MacDonald et al., 2023). Only six non-tailed phage structures have been studied so far: PM2, FLiP and ΦCjT23 which present (*pseudo) p*T = 21 triangulation capsids (Kejzar, Laanto et al., 2022, Laanto, Mantynen et al., 2017, Mantynen, Laanto et al., 2020), and PRD1, Bam35, and PR772 which present *p*T = 25 capsids (Abrescia, Cockburn et al., 2004, Laurinmaki, Huiskonen et al., 2005, Reddy, Carroni et al., 2019). Among them, only FLiP, ΦCjT2, PRD1 and PR772 capsids have been solved at resolution better than 4 Å. In PM2, only the MCP has been solved at high resolution, and the model was combined with lower resolution cryo-electron microscopy (cryo-EM) data of the whole virion (Abrescia, Grimes et al., 2008).

These viruses are diverse in terms of the host cell wall composition, genome types, and living environment. All of them infect Gram-negative bacteria, except Bam35 which infects Gram-positive *Bacillus thuringiensis.* The most common genome form in the DJR lineage is the linear dsDNA genome, shared by PRD1, PR772, and Bam35; PM2 has circular dsDNA genome; while ΦCjT23 and FLiP have circular ssDNA genomes. The environments where these phages were found include soil, plants, intestinal tract, and thermophilic environments, and of course marine environments where they are particularly abundant (Kauffman et al., 2018).

A newly identified non-tailed bacteriophage family, *Autolykiviridae,* has recently been classified in the *Varidnaviria* realm. Autolykiviruses belong to the order *Vinavirales* and are primarily lyic phages that infect marine *Vibrio* species, and are highly abundant in the sea (Kauffman et al., 2018). Notably, non-tailed bacteriophages include the majority of prophages that infect *Vibrio* (Steensen et al., 2024).

Bacteriophage NO16, newly renamed by International Committee on Taxonomy of Viruses (ICTV) binomial nomenclature as *Elsinorevirus NO16* (Turner, Adriaenssens et al., 2025), is a dsDNA temperate vibriophage isolated from *Vibrio anguillarum* (Kalatzis, Carstens et al., 2019). *V. anguillarum* is a pathogenic bacterium producing vibriosis, a mortal haemorrhagic septicemic disease affecting both cultured fish and shellfish (Frans, Michiels et al., 2011). NO16 has recently been classified into a new family named *Asemoviridae*, in the *Vinavirales* order (Kalatzis, Mauritzen et al., 2023, Turner et al., 2025). The global geographic distribution of this viral family and its widespread occurrence as a prophage across >80% of the most prevalent *Vibrio* species, underscores the significance of DJR phages. NO16 and NO16-like phages exhibit a temperate life cycle, enabling their integration into the genome of their *Vibrio* host and spontaneous induction. Lysogenized hosts are more virulent and better biofilm formers compared to their naive counterparts, indicating that NO16 impacts host pathogenicity (Kalatzis et al., 2023).

So far, no structure has been published for any member of this new family, nor any other non-tailed vibriophage, in spite of their abundance and critical ecological role. Here, we report the first structure of a non-tailed vibriophage, one of the simplest members of the DJR lineage. Moreover, we solve the penton/spike symmetry mismatch at the vertex. Using structure and sequence homology search, we build a complete structural atlas of the NO16 infectious cycle.

## Results

### Optimized protocol for NO16 purification

Samples suitable for cryo-EM require a high concentration of particles, homogeneity and structural integrity. We tested previously reported protocols for the propagation and purification of non-tailed phages (Kauffman et al., 2018, Kivela, Kalkkinen et al., 2002, Kivela, Mannisto et al., 1999) and adapted them to obtain a suitable NO16 sample for cryo-EM. The vibriophage purification protocol reported by *Kauffman et al.* (Kauffman et al., 2018), based on ultracentrifugation in a single isopycnic 20-54% iodixanol gradient, yielded a reasonable virus recovery, but was affected by contamination with host membranes (**Fig. S1a-b**). Then, we tested the purification protocol used for the corticovirus PM2, based on an additional sucrose gradient and differential ultracentrifugation step for concentration before the iodixanol gradient (Kivela et al., 1999). Although the double gradient removed the host membranes, the decay of viral titre was pronounced at each step of purification, mainly at the ultracentrifugation concentration step (**Fig. S1a-b**). Additionally, a contaminant co-migrated with the virus in the last gradient, which was identified by LC-MS/MS to be the 60 kDa chaperonin of the host (**Supplementary File 1a**).

To improve viral recovery and remove the 60 kDa contaminant, the concentration step was changed from ultracentrifugation to polyethylene glycol (PEG) precipitation, and an additional iodixanol rate-zonal gradient was designed and included before the last isopycnic iodixanol gradient. This protocol succeeded in obtaining reasonable viral recovery and homogeneous NO16 particles (**Fig. S1a-b**). The main protein band of the purified sample corresponded to the MCP size (29 kDa), confirming the purity of the sample (**Fig. S1c**). This final protocol is detailed in **Figure S1d**.

### Molecular composition of the NO16 virion

Mass spectrometry analysis of the purified sample detected 9 proteins out of the 23 predicted Open Reading Frames (ORFs) in the 10.5 kbp circular dsDNA genome (**Table 1, Supplementary File 1b**) (Kalatzis et al., 2019). Bands consistent with the molecular weight of most of these proteins were observed in the SDS-PAGE of the purified sample (**Fig. S1c**). The structures of all nine proteins detected in the virion were predicted using AlphaFold (**Fig. S2**). The prediction for GP19, the most abundant protein in the virion (**Table 1, Fig. S1c**), included the DJR fold (**Fig. S2**).

**Table 1.**
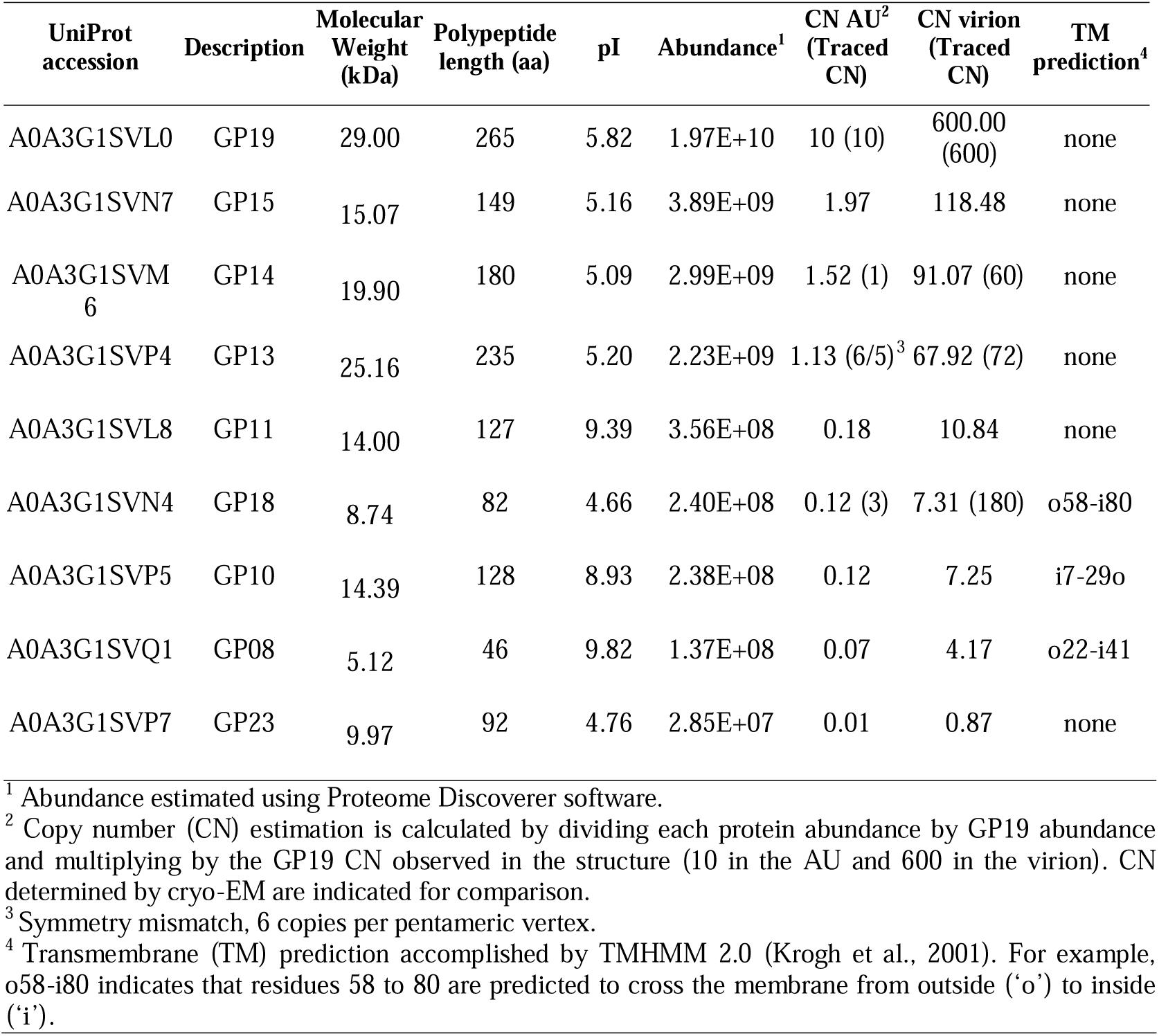
Proteins identified by LC-MS/MS in purified NO16 particles. Ordered by abundance.

### A pT = 21d icosahedral capsid

To solve the structure of bacteriophage NO16, we collected cryo-EM data and applied single particle averaging with icosahedral symmetry (**Table S1, Fig. S3a**). The cryo-EM map at an overall 3.3 Å resolution showed that the NO16 capsid has a triangulation number T = 21d (**Fig. 1a, Fig. S3b-c**). This capsid triangulation number was previously observed in bacteriophages PM2, FLiP and ΦCjT23 (Abrescia et al., 2008, Kejzar et al., 2022, Laanto et al., 2017), and is the smallest reported triangulation number in the DJR lineage. NO16 bacteriophage particles present a diameter of 57 nm facet-to-facet and 70 nm vertex-to-vertex (**Fig. 1a**).

**Figure 1.**
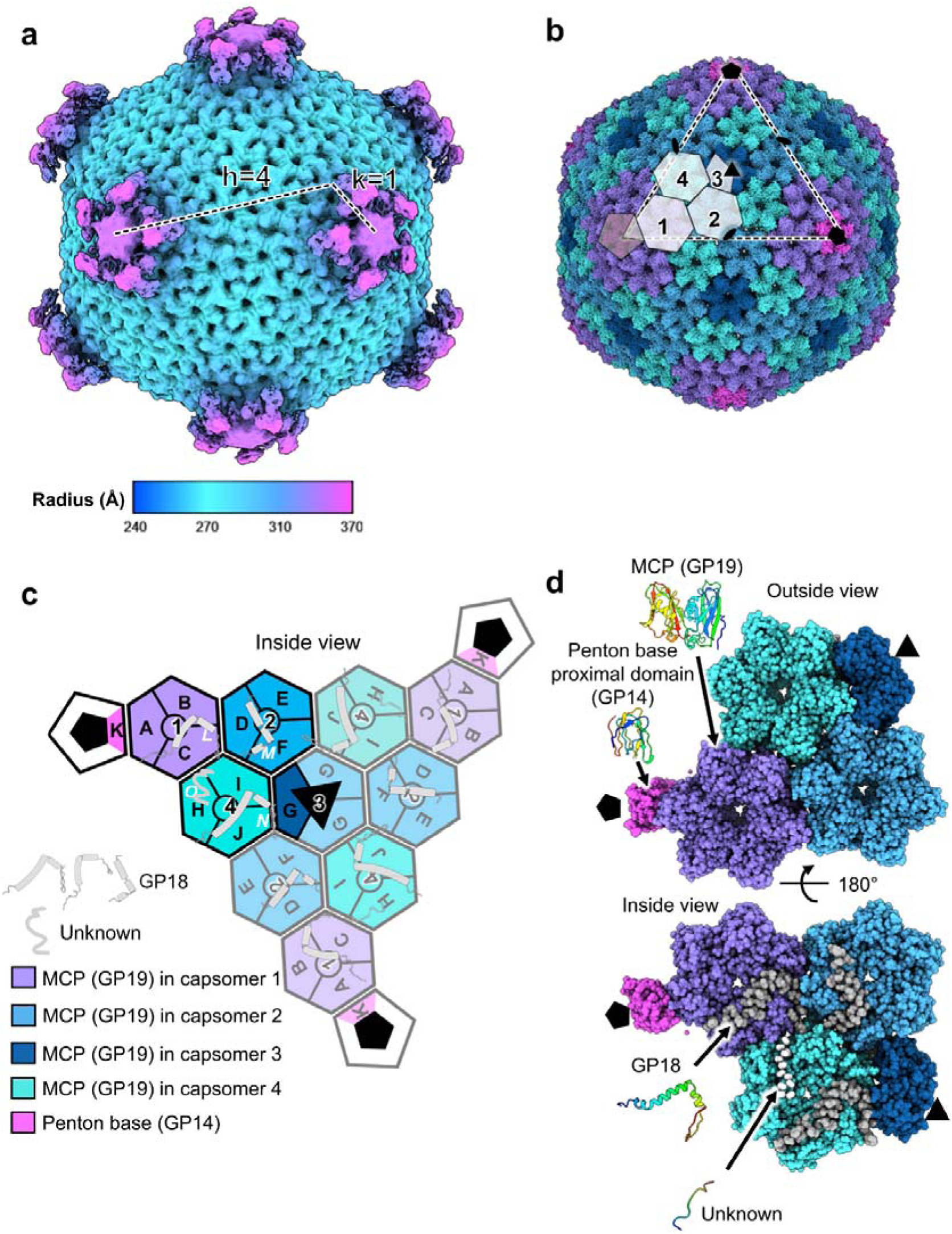
Cryo-EM structure of bacteriophage NO16. (**a**) Cryo-EM map coloured by radius. Dashed black lines define h and k, lattice coordinates or displacement vectors of a pentamer relative to another pentamer used to calculate the triangulation number (T = h^2^ + hk + k^2^) (Caspar & Klug, 1962). (**b**) Molecular models for the proteins traced in the icosahedrally averaged map. The three MCP trimers and one MCP monomer in the asymmetric unit (AU) are numbered 1 to 4. The capsid facet is identified with dashed lines. Icosahedral symmetry axes are indicated with black symbols: fivefold (pentagon), threefold (triangle), and twofold (oval). MCP trimers are coloured according to their position in the AU. Colour legend in (c). (**c**) Schematics showing the organization of one icosahedral facet, as viewed from inside the capsid. One AU is highlighted, and the others are semi-transparent. Letters A-J indicate chain identifiers for MCP monomers. The mCP GP18 chain identifiers are in white. Protein colours and shapes are indicated in the legend. (**d**) Detail of the AU view as seen from outside (top) and inside (bottom). Colour legend in (c). Representative protein structures are rainbow-coloured, from blue (N-terminus) to red (C-terminus).

In the icosahedral averaged map of NO16 capsid, we traced the three proteins (**Table S2-S3**): the MCP (GP19), penton base protein (GP14), and minor capsid protein (mCP) GP18, located on the inner surface of the capsid (**Fig. 1b-d, Supplementary Movie 1**). The asymmetric unit (AU) is composed by 10 MCP monomers, organized into three trimers plus one monomer forming the trimer centred at the icosahedral 3-fold axis, and a monomer of vertex protein (**Fig. 1b**). As MCP form trimeric pseudo-hexamers, the triangulation number of NO16 is defined by pseudo-triangulation (*p*T) = 21d.

### MCP structure: a DJR trimer stabilized by a cation and electrostatic interactions

The MCP (GP19, 265 amino acid (aa) long) presents a conserved DJR fold, with two β-sandwiches separated by a small α-helix at their base and two α-helices on the outer surface (**Fig. 2a**). The NO16 MCP is the shortest one so far reported for the dsDNA *Varidnaviria*, with only the MCP of ssDNA phage ΦCjT23 (P5, 239 aa) being shorter (**Fig. S4a**) (Kejzar et al., 2022). The MCP of its closest relative, the corticovirus PM2, is slightly longer (P2, 296 aa) (**Fig. S4a**) (Abrescia et al., 2008).

**Figure 2.**
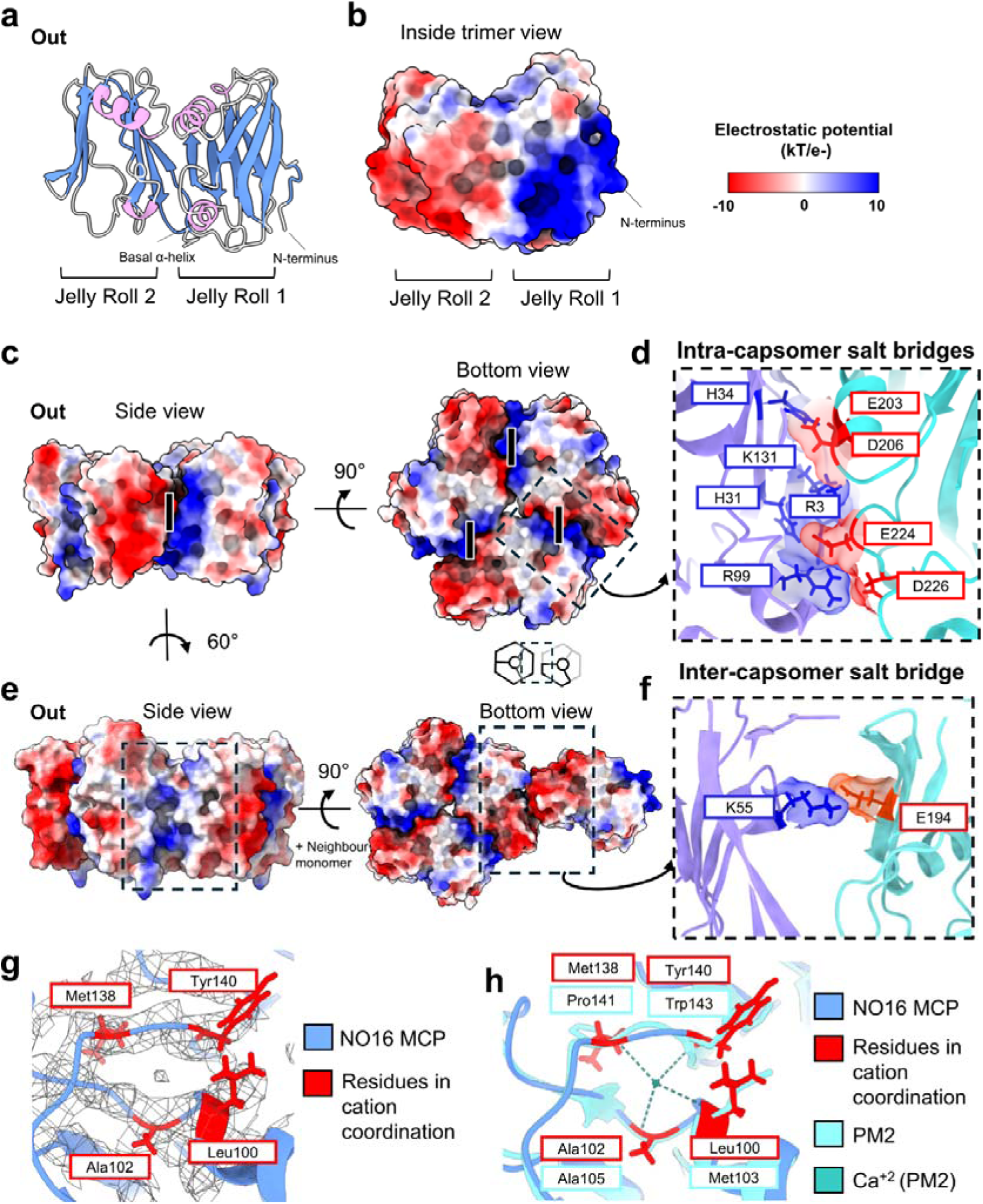
The molecular model for the MCP, GP19. (**a**) Molecular model of the MCP monomer, α-helices are coloured in pink and β-strands in blue. (**b**) MCP monomer surface coloured by electrostatic potential, view from the inside. (**c**) MCP trimer capsomer, from the side (left) and bottom (right), coloured by surface electrostatic potential. I, indicates interface between monomers. The dashed square is zoomed in (d). (**d**) Intra-capsomer surface between monomers, with amino acids taking part in salt bridges highlighted in blue (positively charged amino acids) and red (negatively charged amino acids). (**e**) MCP interface contacting neighbour capsomers indicated in dashed squares (left) and inter-capsomer contact (right), coloured by surface Coulombic electrostatic potential. Schema detailing inter-capsomer contacts. The dashed square is zoomed in (f). (**f**) Inter-capsomer surface between monomers, with amino acids taking part in salt bridges highlighted. (**g**) Cation coordination site at the base of the NO16 MCP with post-processed map threshold at 5.5σ. Residues potentially involved in the interactions are indicated in squares. (**h**) Structural alignment of Ca^+2^ coordinated site of PM2 and NO16 MCP. Residues involved in the interactions are indicated in squares. Model colours indicated in the legend.

The GP19 N-terminal jelly roll (jelly roll 1) is positively charged, while the second (jelly roll 2) is negatively charged (**Fig. 2b**). MCP monomers assemble into trimers to form the capsomers, helped by these strong complementary charges (**Fig. 2c**). Between six and twelve salt bridges can be formed in the NO16 intra-capsomer contact surfaces (**Table S4**), distributed in five unique residue-residue salt bridge interactions (**Fig. 2d**). A similar charge distribution is observed in PM2, but not in FLiP and ΦCjT23 (**Fig. S4b**). Indeed, PM2 MCP is stabilized by up to 10 salt bridges, double the amount detected in ΦCjT23 and FLiP MCP (**Table S5**). In contrast, electrostatic interactions do not play a major role in NO16 inter-capsomer contacts (**Fig. 2e**), with only one residue-residue salt bridge observed (**Fig. 2f, Table S4**).

In the NO16 cryo-EM maps, we observed spherical densities well above noise level (5.5σ) in chains A, E, I and J (**Fig. S4c**), in the same position where a Ca^+2^ ion was observed in PM2 (Abrescia et al., 2008). These densities might correspond to a Ca^+2^ or Mg^+2^ ion since both are present in the buffer (**Fig. 2g, Fig. S4c**). This ion could have a comparable role to that observed in PM2, where it contributes to the stabilisation of the MCP. Indeed, in PM2, treatments with EDTA destabilized the capsid (Kivela et al., 2002) and in the absence of Ca^2+^ the infection was arrested (Cvirkaite-Krupovic, Krupovic et al., 2010). In PM2, the Ca^2+^ ion was in coordination with residues M103, A105, P141 and W143 (Abrescia et al., 2008) (**Fig. 2h**). Structural alignment revealed that in NO16 MCP, residues L100 and Y140 (corresponding to M103 and W143 in PM2) preserve the chemical environment of the Ca^2+^ binding pocket, whereas M138 (P141 in PM2) does not.

### Symmetry mismatched vertex elements probably involved in entry to the host

Penton proteins in DJR viruses are more variable than MCPs. They often present a single jelly roll (SJR) in the penton base (Yutin et al., 2018). In some cases, this SJR is augmented by another protruding SJR, and/or by spike proteins. No penton base nor spike proteins were annotated in the NO16 genome; however, two of the proteins detected in purified virions by mass spectrometry were predicted to have SJR folds: GP13 and GP14 (**Fig. S2**). The proximal N-terminal domain of GP14 (residues 2-89 aa) was identified and traced in the icosahedrally averaged map, forming a SJR buried at the gap left by the peripentonal MCPs (**Fig. 1d**). Density consistent with an additional predicted SJR formed by the distal C-terminal domain was observed, but its lower resolution did not allow tracing the rest of the protein (**Fig. S3c**). Further, additional density formed five petals around the distal part of the vertex. The lower signal of this structural feature suggested the presence of flexible or symmetry mismatched elements at the 5-fold symmetry axis (**Fig. S3c**).

To characterize the five petal densities around the distal domain of GP14 pentamer, we carried out a localized reconstruction analysis of the vertex region (see Methods) (**Fig. S5a-d**). Sub-particle classification revealed that most populated vertex classes included only two opposed petals, one of them with higher density than the other (**Fig. S5c,** red frames). This unexpected symmetry mismatch at the vertex was solved at 3.9 Å resolution in the final map (**Fig. 3a, Fig. S6a-b**). In this local reconstructed map, we traced the GP14 distal domain, which showed that the protein has the conserved penton base structure of a SJR inserted in the capsid shell, and another SJR protruding from the surface of the virus particle (**Fig. 3b-c**). The GP14 proximal and distal domains are connected by a flexible link (**Fig. 3c**).

**Figure 3.**
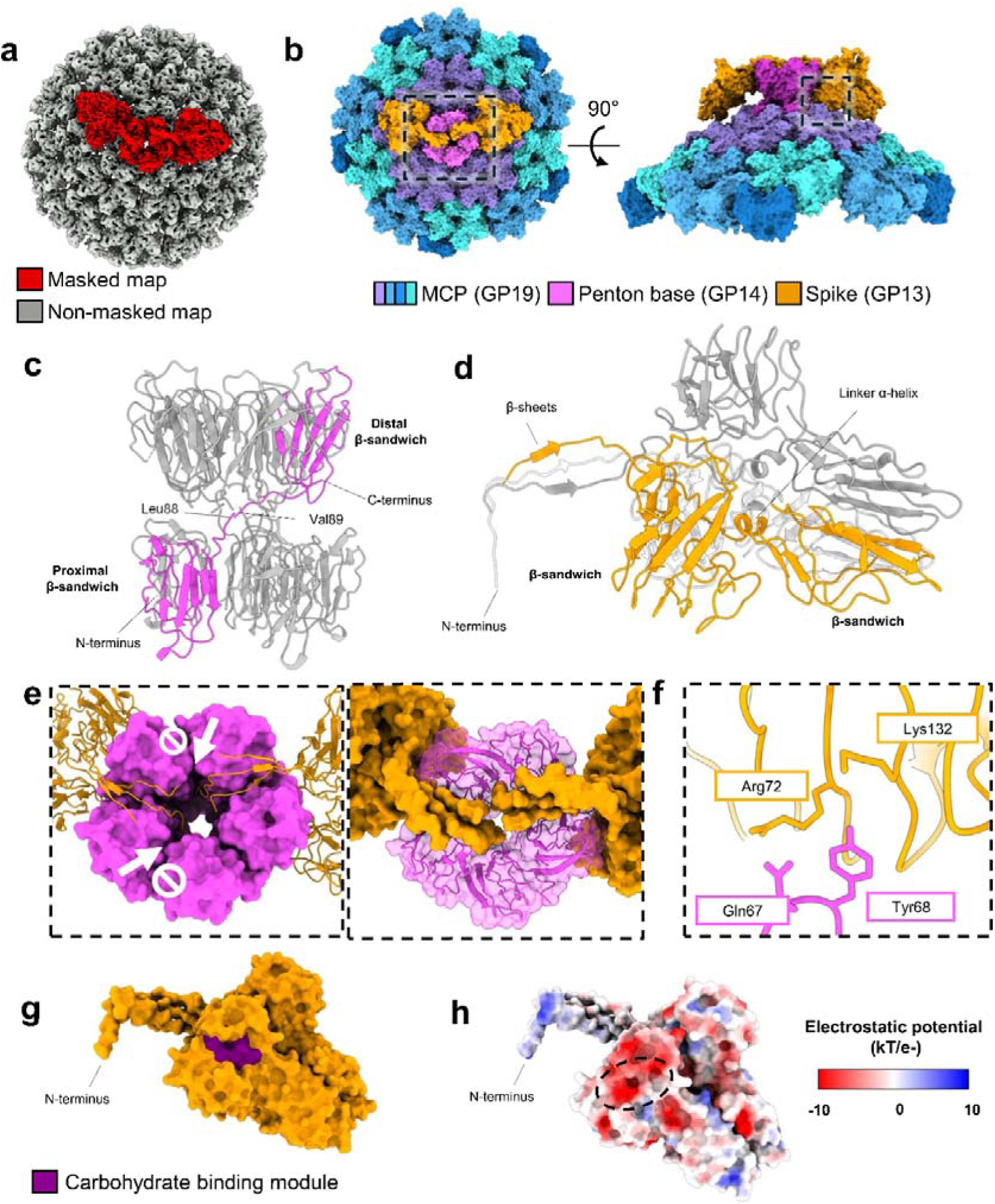
Structure of the vertex region. (**a**) Overlap of local reconstructed maps generated without using any mask (grey) and using a localized mask of the distal penton domain surrounded by two petals (red). (**b**) Molecular model for the vertex region. The dashed square on the left is detailed in (e). The right dashed square is detailed in (f). (**c**) Penton base GP14 structure, with a monomer in pink and the rest of the monomers in grey. (**d**) Spike GP13 trimer structure, a monomer in orange and the rest of monomers in grey and white. (**e**) The N-terminal stalk of GP13 anchoring the spike to the pentamer GP14 central cavity. White arrows and block symbols indicate GP14-GP14 interfaces blocked by GP13 N-terminus. Proteins coloured as in (b). (**f**) Contacts between spike and proximal penton base region. Proteins coloured as in (b). (**g**) Spike trimer highlighting the CBM. (**h**) Spike trimer coloured by electrostatic potential. Dashed oval indicates CBM cleft.

GP13, the second candidate to be the vertex protein, hereafter referred to as spike protein, was identified forming a trimer on the density of each one of the two opposed petal structures (**Fig. 3b,d, Supplementary Movie 1**). The GP13 monomer interacting with the capsid (chain F) was better ordered than distal monomers, which suggests that distal monomers, not stabilized by capsid contacts, display higher flexibility. Therefore, whereas the proximal monomer was fully modelled, flexible distal monomers were partially modelled (**Fig. S6c, Table S3**). The main body of GP13 is composed by two β-sandwich domains, linked by a small α-helix and positioned perpendicularly to each other (**Fig. 3d**).

The two trimeric GP13 spikes bind to the GP14 pentamer in a symmetry-mismatched manner (**Fig. 3b**). In each spike, the N-terminal domains (residues 3 to 20 aa) form an anchoring stalk, composed by three small parallel β-sheets (**Fig. 3d-e**). The two stalks run along GP14 monomer-monomer interfaces in the pentamer. At the pentamer centre, the stalk bends sharply and enters the penton base central cavity (**Fig. 3e**). The N-terminal stalks reach to the neighbouring GP14-GP14 interface, blocking the binding of an adjacent spike (**Fig. 3e**). As a consequence, the second spike must occupy a non-adjacent, opposed position, and addition of a third spike is prevented. This observation aligned with the result of a second localized classification discriminating between presence and absence of spike in a specified position (**Fig. S7**). The class that fixed one spike at position 1 showed two low density spikes in positions 3 and 4, pointing to their partial occupancy, as only one of them could be filled without clashing with the other (**Fig. S6d**). Apart from the N-terminal anchoring, the GP13 monomer positioned most closely to the capsid contacts penton base GP14: Gln67 and Tyr68 of the GP14 N-terminal SJR form hydrogen bonds with GP13 Arg72 and Lys132 (**Fig. 3f, Table S4**). These interactions may contribute to the stabilization of GP13 monomer closer to the capsid surface.

GP13 spikes are parallel to the capsid, differing in orientation from those previously solved, such as PRD1 (Huiskonen, Manole et al., 2007), STIV (Hartman, Eilers et al., 2019) and adenovirus (Martínez, Gallardo et al., 2025) spikes that protrude radially. Homology search with DALI (Holm, 2022) and Foldseek (van Kempen, Kim et al., 2024) showed structural similarity with proteins presenting carbohydrate-binding modules (CBM) (**Supplementary File 2a-b**). DALI indicated that GP13 was most similar to Family 16 CBM S-layer associated multidomain endoglucanase (Bae, Ohene-Adjei et al., 2008) and Foldseek indicated structural similarity to Vip3Bc1 and Vip3Aa protoxin structure showing CBMs (Byrne, Iadanza et al., 2021, Núñez-Ramírez, Huesa et al., 2020) (**Table S6, Supplementary File 2a-b**). The CBM of GP13, located in the N-terminal β-sandwich, displays a curved surface with a cleft (**Fig. 3g**), resembling that of its DALI homologue (Bae et al., 2008), where sugars are thought to interact with polar and acidic residues (**Fig. 3h**). This cleft is solvent-exposed, similarly to protoxin CBMs (Núñez-Ramírez et al., 2020). Notably, the clefts in monomers 2 and 3 did not have enough resolution to trace, consistent with the reported flexible nature of the CBM domains (Núñez-Ramírez et al., 2020) (**Fig. S5c, Table S3**).

Additionally, CLEAN enzymatic function prediction software (Yu et al., 2023) indicated that GP13, even with a low confidence level, could be a glycosylase, a hydrolase that catalyses the cleavage of the glycosidic bond (**Table S6**). Considering that vertex proteins in DJR viruses generally mediate the entry process (Bewley, Springer et al., 1999, Xu, Benson et al., 2003) and that CBM modules are present in tail spike structures in *Caudovirales*, these observations strongly suggest that GP13 could have a role in host wall attachment, recognition and/or hydrolysis.

### A mCP network with little contact with the membrane

We have identified only GP18 as NO16 mCP on the inner side of the capsid (**Fig. 1c-d, 4a**). In the AU, we have traced 3 copies of GP18 beneath capsomers 1 (chain L), 2 (chain M) and 4 (chain N) (**Fig. 1c**). Only the N-terminal domain appears to be ordered (**Table S3**). The C-terminal domain (residues 58 to 80 aa) is predicted to be transmembrane (**Table 1**). However, no density connecting with the membrane was observed (**Fig. S8**). GP18 exhibits complementary charges to the MCP pockets in which it is embedded (**Fig. S9a**) and its binding to the capsomers is energetically favoured, as shown by the negative solvation-free energy (ΔiG), considerably stronger than any other interactions in the asymmetric unit (**Table S4**). This observation suggests that GP18 may act as a hub to help recruit capsomers during assembly. Although they could not be traced due to low resolution, other mCPs seem to be present. For example, chain O was observed next to GP18 in capsomer 4 position (**Fig. 1c-d**), and modelled as a 12 residue-long ‘unknown’ chain (**Table S3**).

Although the icosahedral map did not show additional remnant densities, in local reconstructed maps where the molecular model signal was subtracted (**Fig. S5c**), three main remnant densities appeared (**Fig. 4b**). The main density (D1) is situated beneath the 5-fold axis and mediates contacts with the lipid membrane; a second, small remnant density (D2) is located below capsomer 2 position; and the third and last density (D3) is positioned at the 3-fold axis (**Fig. 4b-c**).

**Figure 4.**
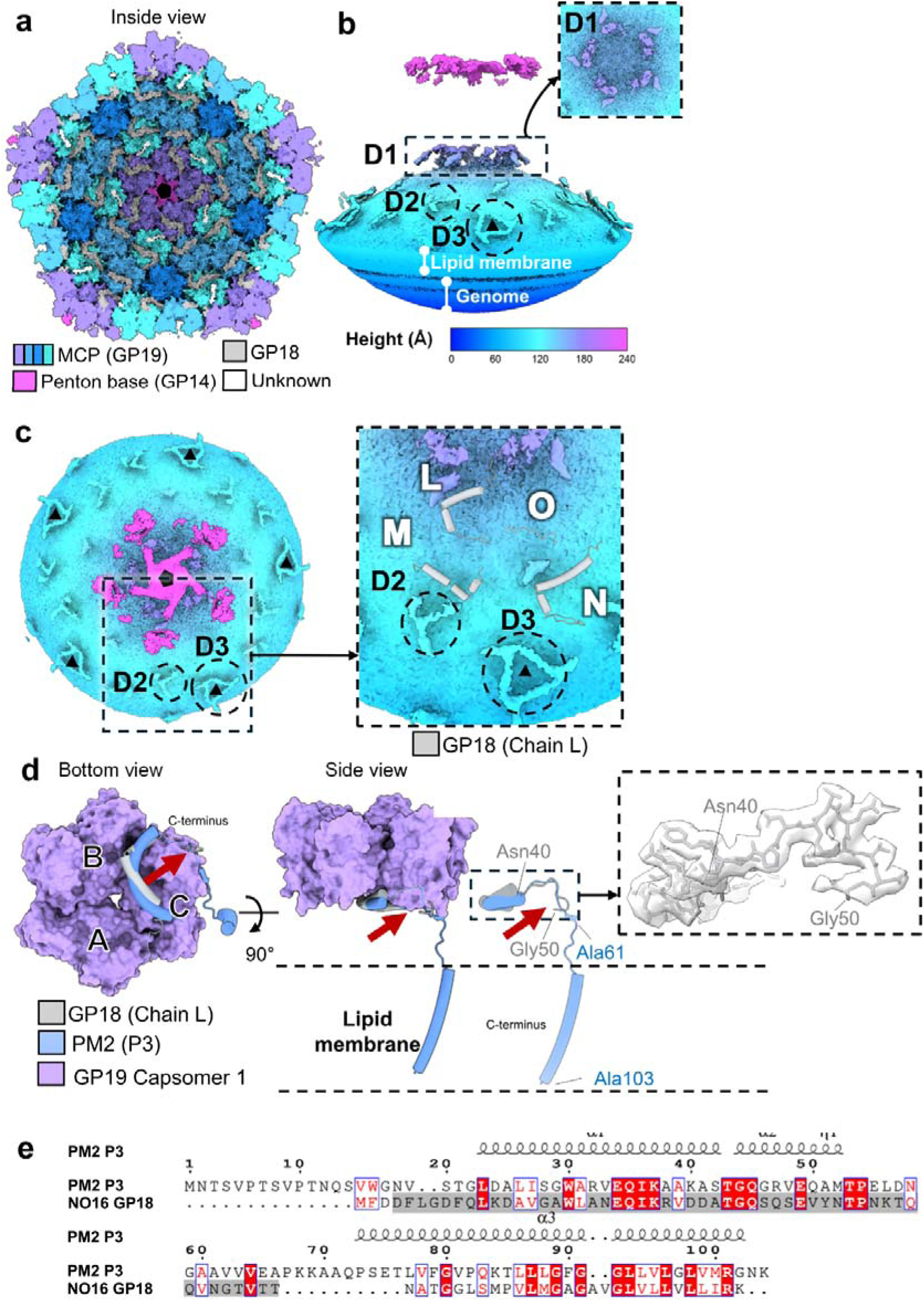
Minor capsid protein organization. (**a**) NO16 inner capsid surface organization viewed along the 5-fold axis. (**b**) Local reconstructed map coloured by heigh showing non traced densities corresponding to mCPs. Dashed panel is zoomed on the top right. (**c**) Local reconstructed map, identical to that in panel (b), rotated –90 degrees along the x-axis. Dashed panel is zoomed on the right. (**d**) GP18 and PM2 P3 protein comparison. Capsomer 1 from inside (left) and lateral (right), GP18 protein and PM2 P3 overlapped. The dashed panel on the right highlights a region of GP18 in chain L, along with its corresponding cryo-EM map density. (**e**) Sequence alignment of PM2 P3 and NO16 GP18 visualized by ENDscript. The traced region of NO16 GP18 (residues 4-55 aa) is highlighted in grey.

To understand GP18 capsid membrane contacts and try to assign an identity to chain O and these remnant densities, we compared with the PM2 structure. GP18 resembles the PM2 mCP P3 in position and sequence identity (39%). Indeed, P3 is a conserved protein among *Vinavirales* non-tailed bacteriophages (Yutin, Rayko et al., 2022). In the PM2 AU, four copies of P3 were traced, with the additional copy being located below capsomer 3. In PM2, the copy of P3 located below capsomer 1 enters the membrane (**Fig. S9b**). In contrast, capsomer 1 GP18 in NO16 does not interact with the lipid membrane, and the pathway followed by the visible part of GP18 indicates a different conformation (**Fig. 4d**) that drastically modifies capsid-membrane contacts. Instead of turning towards the membrane as in PM2, the central region of GP18 turns back towards the inner capsid surface. This conformational change is most evident in the copy near capsomer 1 (**Fig. 4c, red arrows**), but happens in all three copies (**Fig. S9c**). We notice that GP18 is shorter (82 aa) than PM2 P3 (104 aa), which could hinder the ability of the GP18 C-terminal region to reach the lipid membrane, or at least, the interaction could be weaker (**Fig. 4e**).

In PM2, an additional mCP, P6, homologous to NO16 GP10, reinforces the mCP network and contacts the inner lipid membrane as well (**Fig. S9b**). It is located below capsomer 4, at the 2-fold axis and is clearly visible in the central slice of the PM2 cryo-EM map (Huiskonen, Kivela et al., 2004) (**Fig. S8**). In NO16, although GP10 is predicted to be the homologous to P6, it does not correspond to any traced proteins or remnant densities (**Fig. S9d**).

Chain O does not align clearly with any known mCP positions in PM2 (**Fig. S9c**). Additionally, the remnant density D1 is a feature absent in PM2, as there is no electron density below the vertex region (**Fig. S8**). Although the resolution of this density was not enough for identification, taking into account that the protein would necessarily be in the particle and present a transmembrane region, GP10 and GP23 are feasible candidates (**Table 1, Fig. S2**). D2 does not overlap with any of the PM2 mCP either, and its small indistinct shape prevents reliable identification. In contrast, D3 is positioned in the 3-fold axis, precisely where the fourth copy of P3 in PM2 is located (**Fig. S9d**). This positional match strongly suggests that D3 represents an additional copy of GP18.

There is little information on the mCP network for other T = 21 bacteriophages in the lineage. In FLiP, the C-terminal helix of the MCP is believed to interact with the membrane (Laanto et al., 2017), without the need for mCP (**Fig. S8**). In ΦCjT23, although densities contacting the membrane in capsomers 2 and 4 were observed, and the virus encodes P3 and P6 homologues, no mCP was traced (Kejzar et al., 2022). ΦCjT23 may resemble NO16 in that it contains mCP homologous to PM2 and that these mCP do not reach the membrane.

### Completing the NO16 structural atlas

To elucidate putative functions of non-structural proteins present in the virion, as well as proteins transcribed in the host but incorporated into the virus particle, we used AlphaFold structure prediction of protein monomers. Predictions presenting high reliability (pTM larger than 0.75) (**Fig. S2, S7**) were used for structural homology search in DALI (Holm, 2022) and Foldseek (van Kempen et al., 2024). To further support functional predictions, sequence-based CLEAN (Yu, Cui et al., 2023) and InterPro (Blum, Andreeva et al., 2025) analyses were also considered.

### Non-structural proteins present in the virion

Apart from proteins localized in the map, mass spectrometry indicated the presence of another five proteins in the viral particle: GP15, GP08, GP11, GP10 and GP23 (**Table 1**). All these proteins except GP15 have low abundances with an estimation of one or fewer copies per virion (**Table 1**).

Only the GP11 AlphaFold prediction (**Fig. S2**) had high enough reliability to be used for structural homology search. Both DALI and Foldseek detected strong structural homology to the endolysin encoded by *Hafnia* phage Enc34 (**Table S6, Supplementary File 2c-d**). Enc34 is a siphovirus that infects Gram-negative bacteria including the opportunistic *Pseudomonas aeruginosa* PAO1 (Cernooka, Rumnieks et al., 2022). Endolysins are viral peptidoglycan-degrading enzymes that digest the bacterial cell wall (Abdelrahman, Easwaran et al., 2021), usually for release of the virus progeny by lysis of the host. Additionally, CLEAN predicts GP11 to have lysozyme activity (**Table S6**).

### Proteins encoded by NO16 but not detected in the virion

Out of the proteins encoded by NO16 and not detected in the virion (**Fig. S10**), GP03, GP06, GP07, GP16 and GP21 AlphaFold predictions were submitted to homology search, as their predicted models had pTM scores above 0.75, indicating sufficiently high confidence.

DALI indicated that GP03 has slight (Z-score of 9.3) structural homology to *Sulfolobus islandicus* rod-shaped virus 1 (SIRV1) virus protein ORF119, a replication initiation protein, while Foldseek did not identify significant homologues (**Table S6**, **Supplementary File 2e-f**). InterPro supported the possibility that GP03 may function as a replication-associated protein, and CLEAN indicated with low confidence that it can be a DNA ligase. The protein is believed to be part of the rolling-circle synthesis with cleavage and ligase activity (IPR056906).

GP06 is structurally similar to transcriptional repressors (**Table S6**, **Supplementary File 2g-h**). In both DALI and Foldseek searches, λ phage repressor appeared as a candidate. λ phage repressor acts as a genetic switch that enables the phage to transition to lysogeny binding to a lytic promotor (Stayrook, Jaru-Ampornpan et al., 2008). As λ phage repressor is a dimer, GP06 might also form a dimer (Stayrook et al., 2008), and AlphaFold ipTM scores also support this possibility (**Fig. S10**).

Only the N-terminal region of GP07 (residues 1-54) is predicted to be ordered (**Fig. S10**). Homology search of this N-terminal region returned similarities to Anti-clustered regularly interspaced short palindromic repeats (Anti-CRISPR)-associated proteins with both DALI and Foldseek (**Table S6**, **Supplementary File 2i-j**). Phages use anti-CRISPR associated proteins to inhibit host CRISPR-Cas systems, which normally act as transcriptional repressors (Lee, Kim et al., 2022, Liu, Zhang et al., 2021). The prediction indicates that the N-terminal domain of GP07 interacts with the dsDNA and forms a dimer (Liu et al., 2021), suggesting that GP07 might also adopt a dimeric configuration (**Fig. S10**).

For GP16, both DALI and Foldseek indicated homology to acetyltransferases of bacteria (**Table S6, Supplementary File 2k-l**). Acetyltransferases encoded by bacteriophages of the lineage have previously been reported (Yutin et al., 2022).

Finally, GP21 is structurally similar to the genome packaging NTPase of the archaeal virus *Sulfolobus* turreted icosahedral virus 2 (STIV2) of the DJR lineage (**Table S6**, **Supplementary File 2m-n**), which is predicted to be a hexamer (Happonen, Oksanen et al., 2013). AlphaFold ipTM scores for oligomers also support the prediction that GP21 is a hexamer (**Fig. S10**). The genome packaging ATPase is conserved in the realm members as well as in NO16 (Kalatzis et al., 2023, Koonin, Dolja et al., 2022), and is thought to function as a motor for energy-dependent packaging of virus genomes into capsids. InterPro predicts helicase HerA-like domain, found in archaeal helicases and some putative helicases from bacteria. This domain becomes ordered when dsDNA is included in the AlphaFold prediction (**Fig. S10**). CLEAN predicts GP21 to be a site-specific deoxyribonuclease (**Table S6**).

## Discussion

Sample preparation is one of the most significant bottlenecks in cryo-EM and is pronounced in less well-characterized specimens. Production and purification of non-tailed bacteriophages for cryo-EM is reportedly challenging (Bardy et al., 2023). The infection efficiency of NO16 on different populations of *Vibrio anguillarum* varies, even within the same strain (Kalatzis et al., 2023). Moreover, membrane-containing viruses are more sensitive to salinity than non-lipid containing viruses (Kukkaro & Bamford, 2009). Optimization of the purification protocol for NO16 was challenging, and a total of three density gradient steps were necessary to obtain a suitable sample for cryo-EM.

Due to the challenges in purifying non-tailed phages, structural data on them remain scarce, with only four capsid structures solved at near or beyond 4 Å resolution so far (Abrescia et al., 2004, Kejzar et al., 2022, Laanto et al., 2017, Reddy et al., 2019). Moreover, despite achieving high resolution, most of the structures are incomplete since only PRD1 and PR772 structures contain traced mCP proteins (Abrescia et al., 2004, Reddy et al., 2019), while PM2 mCPs were modelled in a cryo-EM map at 8.4 Å resolution (Abrescia et al., 2008).

### One of the simplest capsid in the Varidnaviria realm

The *Varidnaviria* realm contains viruses with genome sizes from 10 kbp to 1,500 kbp and capsid diameters from 60 to 2,500 nm, presenting a wide range of capsid morphologies from T = 21 up to T = 912-1,200 (San Martín & van Raaij, 2018). NO16 presents a T = 21 capsid, which is the smallest observed among realm members, and additionally, its MCP (265 aa) is the shortest reported to date among dsDNA members of the lineage, with only the MCP of the ssDNA phage ΦCjT23 being shorter.

Even though the general organization of NO16 capsid is similar to PM2, especially in the MCP features, the inner organization of the mCP network is simpler. It is widely recognized that cementing proteins are essential for proper viral assembly and, in these viruses, membranes appear to serve as scaffolding platforms that facilitate the recruitment of capsid-forming proteins (Abrescia et al., 2008). However, no ordered membrane-capsid contacts have been visualized in NO16 although tenuous density seemed to connect capsid and membrane near the vertex. This observation indicates that, even if capsid-membrane interactions are needed for assembly, they may not need to be maintained once the particle is formed. This is similar to scaffold protein L1 52/55k, which is processed by adenovirus protease during maturation (Pérez-Berná, Mangel et al., 2014) and to scaffold proteins in *Caudovirales*, which are also removed during the transition from procapsid to mature capsids (Prevelige & Fane, 2012).

Given the structural features of the MCP and the mCP network, NO16 is simpler than PM2, and therefore, the simplest dsDNA virus of the DJR lineage and *Varidnaviria* realm, with the ssDNA virus ØCjT23 being the simplest one of the realm.

### Vertex symmetry mismatch, intrinsic of receptor binding proteins

It is quite common for spike proteins to show symmetry mismatches and flexibility, as in PRD1 (Huiskonen et al., 2007, Merckel, Huiskonen et al., 2005) or adenovirus (Chiu, Wu et al., 2001, Martínez et al., 2025, Wu, Pache et al., 2003). In NO16, we have found two trimers of spike protein GP13 forming the petals, assembled in a symmetry-mismatched manner to the GP14 pentamer. In previous works in *Varidnaviria* bacteriophages, attempts at local reconstruction failed to obtain enough resolution to trace the spike proteins (Kejzar et al., 2022, Reddy et al., 2019). Apart from tailed phages and their herpesvirus relatives, where prominent structural features and strong cryo-EM signals have enabled the successful resolution of symmetry mismatches, this represents, to the best of our knowledge, the first time in which local reconstruction solved the symmetry-mismatched vertex region at enough resolution to trace the spike protein.

The spike protein GP13 presents a CBM and a putative glycosylase activity. Archaeal DJR virus STIV also exhibits a CBM in the spike protein forming the turret (Hartman et al., 2019). It has been proposed that most of the viral proteins were recruited from cellular ancestral proteins, and indeed the β-sandwich structure is found in cellular proteins, most of which bind carbohydrates and some are carbohydrate-active enzymes (Koonin et al., 2022, Krupovic & Koonin, 2017). In this particular case, the CBM module present on the ancestral protein could have been maintained through exaptation to play a role in the attachment and/or entry to the host. In bacteriophage Bam35, which infects Gram-positive bacteria, a sugar unit of the peptidoglycan, *N*-acetyl-muramic acid was identified as a substrate of the Bam35 receptor (Gaidelyte, Cvirkaite-Krupovic et al., 2006).

The presence of three monomers of GP13 in each spike would be advantageous by increasing the affinity for polysaccharides. Moreover, the two trimers located diametrically opposed in each vertex may enhance the search space for host receptors. Additionally, blurry densities visualized on top of the vertex (**Fig. S6e**), might indicate mobility of spikes allowing to explore the surrounding space and facilitate receptor binding (**Fig. 5**).

**Figure 5.**
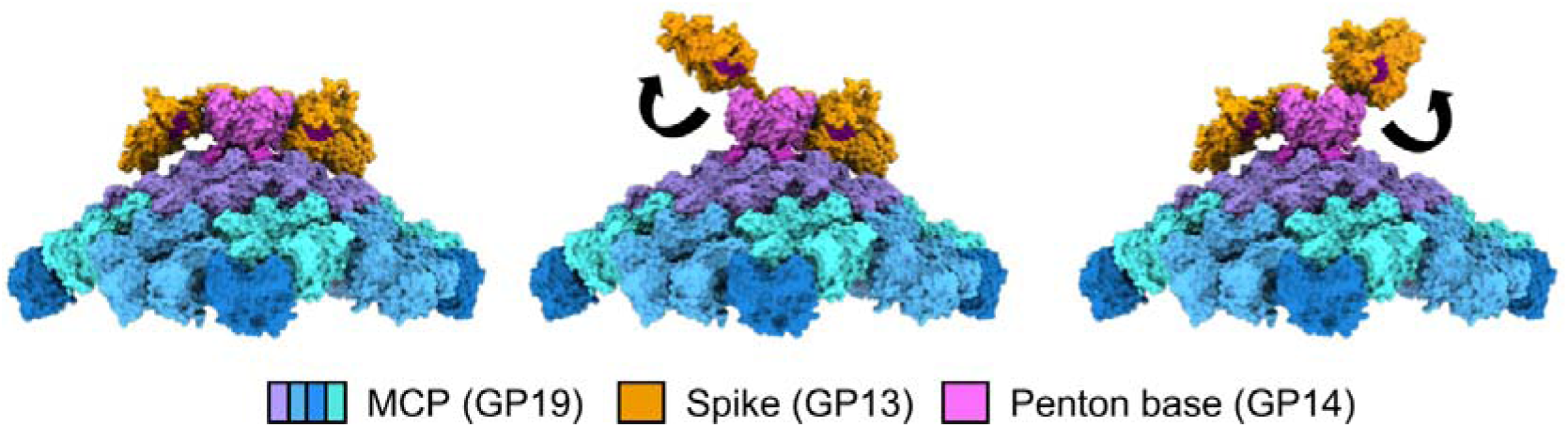
Putative model for GP13 spike mobility for receptor binding.

### A hypothetical model for the NO16 infectious cycle

Putting together the structure of the NO16 particle and the predictions for the non-structural proteins, we propose a hypothetical model for the NO16 infectious cycle (**Fig. 6**). As a first step in the infection, NO16 must cross the outer membrane, the peptidoglycan layer and the inner membrane. Considering vertex proteins usually mediate entry (Bewley et al., 1999, Xu et al., 2003), the spike protein GP13, with its CBM module and putative glycosylase activity, could be involved in both attachment and entry (**Fig. 6, 1-2**). Additionally, putative endolysin GP11, with predicted lysozyme activity, is present in the viral particle. Although endolysins have been associated mostly with the end of the replication cycle, the presence of this activity in the viral particle raises the intriguing possibility that GP11 could play a role in degradation of the peptidoglycan layer during NO16 entry (**Fig. 6, 2**). Similarly, bacteriophage PRD1 particles carry a membrane-associated transglycosylase that appears to play a role during the early stages of infection (Rydman & Bamford, 2000). In related bacteriophages of the *Vinavirales* order, which include NO16 family *Asemoviridae, Autolykiviridae, Corticoviridae, Mestraviridae* as well as *Parnassusviridae*, peptidoglycan hydrolysers or endolysins have been described to be conserved (Yutin et al., 2022). We propose that GP13 and GP11 could be potential candidates as lytic enzymes against *V. anguillarum* infections, which cause significant losses in fish farms (Frans et al., 2011).

**Figure 6.**
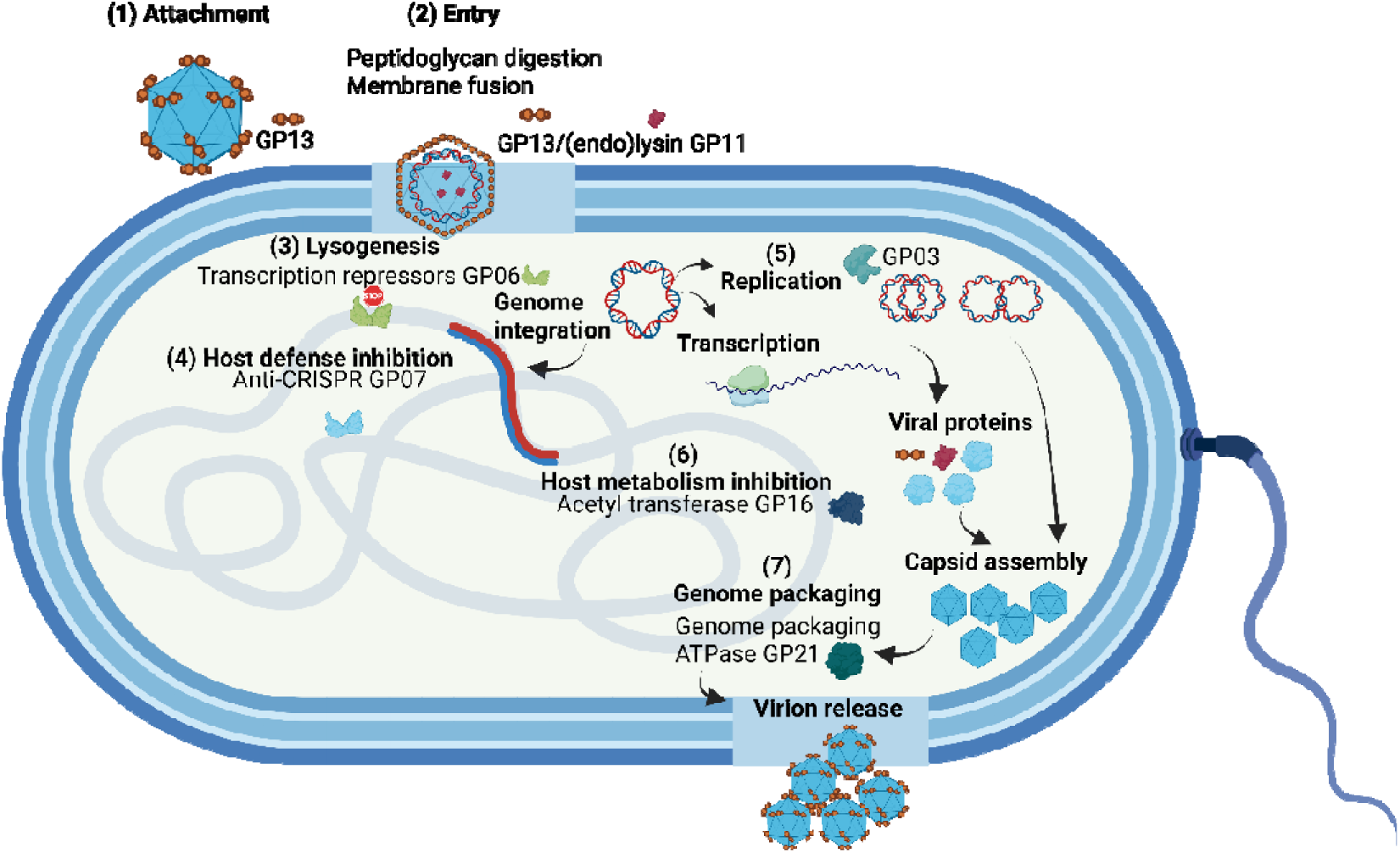
Hypothetical model for the NO16 infectious cycle. (1) First, NO16 attaches to the host cell wall via the CBM of GP13. (2) Entry occurs by carbohydrate degrading capacity of GP13 and (endo)lysin GP11, and most probably via membrane fusion. (3) Once the genome is released into the host cell, it integrates into the host genome and expression of transcription factor GP06 regulates the lytic/lysogenesis switch. (4) GP07 anti-CRISPR associated protein could inhibit host defence. (5) Replication via the rolling circle mechanism proceeds with participation of GP03. (6) GP16 acetyltransferase modulates host proteins by acetylating them. (7) The viral structural proteins assemble around host membranes, and the genome is packaged with the help of ATPase GP21. Created by Biorender.com.

Two entry mechanisms have been proposed for *Varidnaviria* bacteriophages: by forming a tube-like structure facilitating genome delivery in the linear-genome phage PRD1 (Peralta, Gil-Carton et al., 2013), and by membrane fusion in the circular-genome phage PM2 (Cvirkaite-Krupovic et al., 2010). Since we have not observed any tube-like structure in the cryo-EM micrographs and the genome of NO16 is circular dsDNA as in PM2, we hypothesise that NO16 entry could be by membrane fusion (**Fig. 6, 2**), similarly to that proposed for PM2 (Cvirkaite-Krupovic et al., 2010).

Once inside the host cell, the NO16 genome integrates into the host genome through a site-specific recombination mechanism in a lysogenization process that is favoured at high host cell concentrations, suggesting that the lysis-lysogeny switch is likely regulated by host density-dependent mechanisms (e.g. quorum sensing) (Kalatzis et al., 2023). GP06 and GP07 expression was enhanced during conditions that favoured lysogeny (Kalatzis et al., 2023), supporting their role in regulating prophage induction/integrations dynamics. Further, GP06 is a structural homolog of λ phage repressors which bind to lytic promoters (Stayrook et al., 2008) (**Fig. 6, 4**). Similarly, in non-tailed Jorvik phage gp1 plays a role in lysis-lysogeny decision (Bardy et al., 2023). For GP07, we hypothesize an anti-CRISPR associated function, which could block bacteria defence systems against infections (**Fig. 6, 4**). A similar function has been attributed to STIV B116 protein (Overton, Manuel et al., 2023).

Since GP03 is homologous to PM2 replication initiation protein and the NO16 genome is circular dsDNA, it seems likely that NO16 replication occurs via the rolling circle mechanism, with GP03 involved in the initiation process near the cytoplasmic membrane (Brewer, 1978, Espejo, Canelo et al., 1971, Mannisto, Kivela et al., 1999) (**Fig. 6, 5**). According to the structural homology of the replication proteins, the phage only codes for the replication initiation protein, and the other necessary proteins should be hijacked from the host. Mass spectrometry analysis of the purified virion revealed the presence of two host histone-like DNA-binding proteins (UniProt: A0A191W6X7 and A0A191W301) and the host DNA topoisomerase (UniProt: A0A1Y0NSD3), with higher abundance than some mCPs (**Supplementary File 1b**). However, it remains unclear whether these proteins are components of the virion or were purified as contaminants. The replication mechanism of the circular dsDNA genomes in the DJR lineage is not well understood yet, since it is a different genome replication process than the common mechanism in the realm driven by the conserved type B DNA polymerase (PolB) (Kauffman et al., 2018, Krupovic & Bamford, 2007). Notably, PolB is not encoded by any of the circular dsDNA or ssDNA non-tailed phages ΦCjT23, FLiP, PM2,and NO16. Moreover, the replication system components within bacteriophages of *Vinavirales* order are frequently replaced (Yutin et al., 2022), likely to adapt to different types of genome (circular ssDNA, linear dsDNA and circular dsDNA).

GP16, a putative acetyltransferase, could be altering host metabolism during infection (**Fig. 6, 6**) (Christensen, Xie et al., 2019). Acetyltransferases can, by adding the acetyl group, neutralize protein charge and therefore, change protein function dramatically (Christensen et al., 2019). They also have a specific role in histone acetylation in eukaryotic cells (Bannister & Kouzarides, 2011). Members of the GCN5-related *N-*acetyltransferases (GNAT) have been detected in multiple *Vinavirales* genomes, although not in PM2 (Yutin et al., 2022). Certain GNATs are associated with histone acetyltransferase activity (Favrot, Blanchard et al., 2016). There is recent evidence that bacteria encode proteins with hypothetical histone domains (Hocher, Laursen et al., 2023), and *V. anguillarum* proteins have been annotated as histone-like DNA-binding proteins (UniProt: A0A191W6X7 and A0A191W301). GP16 could, therefore, have a role in inhibiting bacterial host metabolism and/or host chromatin remodelling.

In the late stage of infectious cycle, the viral proteins assemble around host membranes into capsids and the genome is incorporated with the help of genome packaging ATPase GP21 (**Fig. 6, 7**) (Happonen et al., 2013). Viral genomes are packaged into capsids by one of two mechanisms: concerted, where the protein shell is built around the genome (Condezo & San Martín, 2017, Garber, Seidman et al., 1980, Gómez-Gónzalez, Burkhardt et al., 2024); or sequential, where the genome is pumped into a preformed empty capsid (Cuervo, Dauden et al., 2013). The genome packaging ATPase is conserved in the realm and plays a role in sequential packaging mechanisms (Hong, Oksanen et al., 2014) as well as in concerted packaging (Condezo & San Martín, 2017). Therefore, the presence of genome packaging ATPase in NO16 does not clarify the packaging mechanism.

In conclusion, this study elucidates new high-resolution data among the understudied *Varidnaviria* bacteriophages. This is the first structure of a non-tailed vibriophage, and of a phage in the newly defined *Asemoviridae* family. Notably, it also reveals the first symmetry mismatched vertex complex solved at high resolution among viruses of the lineage (**Supplementary Movie 1**). To further complete the structural atlas of NO16, and considering 70% of structural homologs that could be identified between viral and cellular proteins have no detectable sequence homology (Lasso, Honig et al., 2021), we employed tertiary structure prediction tools and structural-based homology programs to assign the function of some viral proteins within the viral cycle. While these findings offer valuable insights, further experimental validation is necessary to confirm the proposed infection model. Studying the structural and functional details of viruses infecting bacterial pathogens, such as *Vibrionaceae*, is essential for the exploration of new therapeutic potentials.

## Materials and Methods

### Virus and bacterial strains

Bacteriophage NO16 (GenBank: MH730557) and its host *Vibrio anguillarum* strain A023 (GenBank: JAHGUX000000000) were available in the Prof Middelboe laboratory (Kalatzis et al., 2019). NO16 lysate was stored at 4°C and stock of the host was stored at −180°C with 20% glycerol.

### Host culture

*V. anguillarum* was cultured in Marine Broth medium (MB; 0.5% tryptone, 0.1% yeast extract, and 2% sea salts, Himedia M1942-500G), incubated at 25°C in agitation. *V. anguillarum* colonies in MB 1.5% agar plates were stored at room temperature and inoculated into liquid medium for growth. Culture growth was monitored by optical density values at 600 nm wavelength (OD_600_).

### NO16 propagation and purification

*V. anguillarum* A023 culture was infected at OD_600_ = 0.15 (1×10^8^ cells/ml) with NO16 bacteriophage lysate at MOI = 0.1, and incubated at 25°C in agitation for 3 hours approximately until detection of complete lysis (OD_600_ ≤ 0.05). Cell debris was removed by centrifugation at 18,800 xg, 20 minutes, 4°C and the supernatant was filtered through 0.45 µm filters (MF-Millipore™, HAWP304F0). The virus was concentrated by precipitation in 10% (w/v) PEG 8000 overnight at 4°C and diluted in PM2 buffer with magnesium (PM2-Mg buffer: 20 mM Tris-HCl pH 7.2, 100 mM NaCl, 5 mM CaCl_2_, 8 mM MgSO_4_ x 7 H_2_O) (Kivela et al., 1999).

Purification of the bacteriophage was performed by ultracentrifugation in three subsequent gradients. First, the precipitated virus was loaded onto a linear 5-20% sucrose gradient diluted in PM2-Mg buffer for 1.5 hours at 24,000 rpm and 15°C (SW41 rotor). The pellet containing membranous contaminants was removed and the upper fractions were concentrated again with 10% PEG 8000. The precipitate was loaded onto a linear gradient of 8-16% iodixanol diluted in PM2-Mg buffer (OptiPrep™) with a 36% iodixanol cushion, and ultracentrifuged at 15°C, 30,000 rpm, 35 minutes (SW55 rotor). The upper part of the cushion (approximately 1.5 ml) was loaded onto a 9-layer iodixanol gradient (21-24-27-30-33-36-42-48-54% iodixanol diluted in PM2-Mg buffer), and centrifuged for a minimum of 10 hours at 35,000 rpm and 18°C (SW41 rotor). The virus bands (running at approximately 30% iodixanol) were collected and stored at 4°C until further use. Prior to experiments, the purified NO16 virus sample was dialysed against PM2-Mg buffer.

### Infectious titre determination

Double-layer agar assays were performed on M6 plates. For the bottom agar, 4 ml of MB medium with 1.5% agar were stirred in each well and left to harden. A volume of 1 ml of top agar (0.4% agar and 1% sea salt) was mixed with 100 µl of OD_600_ ∼ 0.15 *V. anguillarum* culture and 10 µl of serial dilutions of the bacteriophage. Plates were stored at room temperature overnight and lysis plaques were counted to estimate plaque-formation units per millilitre (PFU/ml).

### Negative-staining electron microscopy

To assess the homogeneity and structural integrity of purified virus samples, they were dialysed against PM2-Mg buffer for one hour at 4°C, and imaged by negative staining electron microscopy. In-house collodion-carbon coated copper grids (Gilder G400-C3) were glow discharged for 15 seconds at 25 mA in a vacuum chamber to overcome the hydrophobicity of the carbon surface. Subsequently, samples were incubated on the grid for 5 minutes, stained for 30 seconds with 3 µl 2% uranyl acetate, and blotted to remove the excess of liquid. Micrographs were obtained with a 120 kV JEOL JEM 1011 TEM at the CNB-CSIC Electron Microscopy Facility.

### Denaturing protein electrophoresis (SDS-PAGE)

Samples extracted from the gradients were mixed with 3x sample buffer (1% SDS, 1% β-mercaptoethanol, 10% glycerol, 50 mM Tris-HCl pH 6.8, 1.6% bromophenol blue) and heated at 95°C for 10 minutes. Proteins were separated by SDS-PAGE using 4-20% acrylamide gradient gels (Bio-Rad, 4568093). The electrophoresis was carried out for one hour at 90 V.

After electrophoresis, gels were fixed for at least one hour (up to overnight) in 40% ethanol and 10% acetic acid, then sensitized in 30% ethanol, 125 mM sodium thiosulfate, and 800 mM sodium acetate for 30 minutes, and washed three times with milli-Q water for 3 minutes. Afterwards, gels were stained with 0.25% silver nitrate for 20 minutes and washed twice for 30 seconds. Finally, gels were soaked in 240 mM sodium acetate and 0.015% formaldehyde until protein bands were distinguishable. The reaction was stopped by 40 mM ethylenediaminetetraacetic acid (EDTA) disodium salt dihydrate (EDTA-Na_2_ x 2H_2_O).

### Mass spectrometry

#### In gel digestion

To identify the NO16 purification contaminants after SDS-PAGE, the band at 60 kDa was excised, cut into 1 mm^3^ pieces and placed in a 96-well plate for automated trypsin digestion in an OT2 robot (Opentrons) as described elsewhere (Shevchenko, Wilm et al., 1996) with minor modifications. Briefly, gel pieces were first washed with 50 mM ammonium bicarbonate and then with acetonitrile (ACN) before simultaneous reduction and alkylation with 5 mM tris(2-carboxyethyl)phosphine hydrochloride (TCEP) and 10 mM chloroacetamide (CAA) in 50 mM ammonium bicarbonate. The gel pieces were then rinsed, first with 50 mM ammonium bicarbonate and then with ACN. Finally, they were dried at 40°C. Proteomics grade trypsin (Pierce) was added at a final concentration of 16 ng/μl in 25% ACN, 50 mM ammonium bicarbonate and digestion was allowed to proceed at 37°C for 4 hours. Peptides were extracted by incubation with 50% ACN, 0.5% trifluoroacetic acid (TFA) for 25 minutes. They were then speed-vac dried and stored at −20°C until liquid chromatography coupled to tandem mass spectrometry (LC-MS/MS) analyses.

#### Liquid S-Trap^TM^ digestion

The purified NO16 sample was diluted until 5% sodium dodecyl sulphate (SDS) and 25 mM triethylammonium bicarbonate (TEAB). The sample was reduced and alkylated by addition of 5 mM TCEP and 10 mM CAA for 30 minutes at 60°C. Protein digestion in the S-Trap filter (Protifi, Huntington, NY, USA) was performed following the manufacturer’s instructions with minor changes (Ciordia, Alvarez-Sola et al., 2021). The sample was digested at 37°C overnight using a trypsin:protein ratio of 1:15 and desalted with a StageTip C18 prior to analysis by LC-MS/MS.

#### LC-MS/MS Analysis

Approximately 400 ng of peptides were dissolved in formic acid and subjected to LC-MS/MS analysis using an Ultimate 3000 nanoHPLC (Thermo Fisher Scientific) coupled online to an Orbitrap Exploris 240 mass spectrometer (Thermo Fisher Scientific). The HPLC was equipped with an Easy[spray PepMap C18 analytical column (50[cm[×[75 μm; Thermo Fisher Scientific). Solvents A and B were, respectively, 0.1% formic acid and 0.1% formic acid in 80% acetonitrile. Peptides were separated with a 47 min gradient ranging from 4% to 95% B (60 minutes of total run time) at 50°C and a flow rate of 250[nl/min.

Data acquisition was performed using a data-dependent top 20 method in positive ionization mode. Survey scans were acquired at a resolution of 60,000 (FWHM). The top 20 most intense signals from each MS1 scan were selected for fragmentation by Higher-energy Collisional Dissociation (HCD) 30%. Resolution for MS2 spectra was set to 15,000 (FWHM). Precursor isolation was performed with a window of 1 m/z.

#### Database Search and Protein Identification

Raw files were processed with Proteome Discoverer (Orsburn, 2021) (version 2.5.0.400; Thermo Fisher Scientific). MS2 spectra were searched with MASCOT (Koenig, Menze et al., 2008) (version 2.8.0; Matrix Science), MSFragger (Kong, Leprevost et al., 2017) and Sequest against a database with the reference proteomes of bacteriophage NO16 (UniProt: UP000276974), its host, *Vibrio anguillarum* (UniProt: UP000197971), and a set of common contaminant sequences. Identifications were filtered at a false discovery rate (FDR) < 1% at the protein, peptide, and peptide-spectrum match (PSM) levels. Mass spectrometry analyses were completed at the CNB-CSIC Proteomic Facility.

For the purified NO16 sample, copy number estimation was calculated by dividing the abundance of each protein by that of GP19 (the MCP) and multiplying by the number of GP19 copies present in the virion. The transmembrane prediction of each protein was accomplished by TMHMM 2.0 (https://services.healthtech.dtu.dk/services/TMHMM-2.0/) (Krogh, Larsson et al., 2001). The result tables are included as **Supplementary File 1.**

### Cryo-EM sample preparation and data collection

Purified virus samples (1.5×10^11^ PFU/ml) were dialysed against PM2-Mg buffer for one hour at 4°C and deposited in glow-discharged (1-minute, 25 mA) Quantifoil R1.2/1.3 Copper/Rhodium (Cu/Rh) grids. Three consecutive incubations on the grid with 3 µl drops of the sample followed by blotting were performed to improve the amount of particles lying in the carbon holes (Snijder, Borst et al., 2017). Excess liquid was blotted and grids were vitrified using an FEI Vitrobot. Grids were examined in a 200 kV Talos Arctica microscope at the CNB-CSIC cryo-EM facility, to assess particle concentration, ice quality and overall suitability of the sample. Cryo-EM data were collected at the Diamond Light Source Facility (eBIC, Oxford, United Kingdom) using a 300 kV Titan Krios microscope equipped with a Bioquantum K3 camera at a nominal pixel size of 1.34 Å at 64,000 x. Additional vitrification conditions and imaging parameters are summarized in **Table S1**.

### Cryo-EM single particle averaging with icosahedral symmetry

All image processing was performed within the Scipion 3 framework (de la Rosa-Trevin, Quintana et al., 2016). Frame alignment was carried out with using MotionCor2 (Zheng, Palovcak et al., 2017). The contrast transfer function (CTF) was estimated using CTFFIND4 (Rohou & Grigorieff, 2015). Particles were semi-automatically picked using Xmipp3 (de la Rosa-Trevin, Oton et al., 2013) and extracted in 900 x 900 px boxes. 2D and 3D classifications were conducted in RELION (Scheres, 2012). Icosahedral symmetry was enforced for 3D classification and refinement. The best 3D class was further processed by RELION 3D auto-refine followed by CTF refinement (Zivanov, Nakane et al., 2018). The corrections were applied in the following order: beam tilt refinement, anisotropic magnification, and per particle defocus correction. Finally, 3D auto-refine was run again followed by resolution estimation (Fourier Shell Correlation (FSC) at 0.143 threshold) and high frequency boosting (B-factor, RELION post-process). The final map yielded a 3.3 Å resolution (**Fig. S3b**) (Russo & Henderson, 2018). Additional image processing, reconstruction and refinement parameters are summarized in **Table S1**. Local resolution was calculated with MonoRes and depicted in ChimeraX (Vilas, Gomez-Blanco et al., 2018).

### Local reconstruction of vertex regions

Vertex sub-particles centred on the 5-fold axis at the capsid radius were extracted with Localrec (Abrishami, Ilca et al., 2021) in 320 x 320 px images (**Fig. S5a**). 60 sub-particles were generated from each virus particle and all downstream image processing was carried out without enforcing any symmetry. Due to uncertainty in the handedness, the extraction was performed twice by changing the sign of the defocus correction. Both sets of sub-particles were reconstructed in parallel, and the subset yielding the higher resolution was selected for further processing. Next, extracted particles were realigned with CryoSPARC local refinement (Punjani, Rubinstein et al., 2017). To enhance the signal of the five upper petals, we subtracted the capsid projection signal with RELION (Scheres, 2012) in two stages. First, a mask based on the structure of the AU traced in the icosahedral map was used to obtain an initial model of the penton base GP14 distal domain and properly fit the AlphaFold prediction of this domain. Second, a mask based on both the structure of the AU traced in the icosahedral map and the AlphaFold prediction of the penton base (GP14) distal domain was used to isolate the density corresponding to the five petals surrounding the penton base (**Fig. S5b**). Subtracted particles were then classified in 10 classes with CryoSPARC (**Fig. S5c**). The sub-particles of the five classes showing at least one petal with high signal to noise ratio were selected and merged (**Fig. S5c, red box**). To improve map resolution, we re-aligned sub-particles rotating them by a multiple of 2П/5 rad to average the highest petal signal of every class (**Fig. S5c**). The new pool of re-aligned sub-particles was replaced by identical aligned sub-particles without capsid subtraction for further RELION refinement. Redundant particles were removed before calculation of final maps (**Fig. S5d**). Two refined maps were generated, with or without using any mask and using a localized mask of the distal penton domain surrounded by two petals (**Fig. S5d**), at 3.9 Å and 3.7 Å resolution, respectively. More details of the workflow are summarized in **Fig. S5** and in a *json* file (**Supplementary File 3**). Additional local reconstructed maps were generated focusing the refinement in one petal instead of at the 5-fold axis using a workflow summarized in **Fig. S7**.

### Model building and analysis

Model building was completed within the Scipion 3 framework (Martínez, Jiménez-Moreno et al., 2020). The initial model for GP19 and GP14 was predicted using AlphaFold 2 (Jumper, Evans et al., 2021). ChimeraX was used to perform a rigid fitting of each chain initial model into the sharpened map (Meng, Goddard et al., 2023). Next, the fitted model of each chain was refined using Coot (Emsley, Lohkamp et al., 2010) and Phenix real-space refine (Liebschner, Afonine et al., 2019). To identify additional unmodeled densities, we generated remnant maps with ChimeraX by creating a map from the traced model (*molmap* command) and subtracting it from the main cryo-EM map. ModelAngelo (Jamali, Kall et al., 2024) was used to generate initial molecular models in the remnant map. In the first ModelAngelo search, no sequence was provided, and results were submitted to BLAST (https://blast.ncbi.nlm.nih.gov/Blast.cgi) against the NO16 proteome. When hits had significant e-value, a second ModelAngelo search was completed with the protein sequence of the hit. The resulting molecular models were refined using Coot and Phenix real-space refine.

Map and model visualization, including Coulombic electrostatic potential colouring, were carried out with ChimeraX (Pettersen, Goddard et al., 2021). For the structural alignment of the models, ChimeraX *matchmaker* command was used (Meng et al., 2023). Interactions between traced proteins were analysed using PDBePISA (Krissinel & Henrick, 2007). Sequence alignments between NO16 GP18 and PM2 P3 were carried out with Clustal Omega (Madeira, Madhusoodanan et al., 2024), and visualized using ENDscript (Robert & Gouet, 2014).

### Tertiary structure prediction, structural homolog searches and function prediction

The structure of NO16 proteins not traced in the cryo-EM map was predicted using AlphaFold 3 (https://alphafoldserver.com/) (Abramson, Adler et al., 2024). The models were assessed based on predicted Template Modelling (pTM) score which evaluates the accuracy of a protein structure prediction. When multimers or additional cofactors were included in the prediction, the interface predicted Template Modelling (ipTM) score was used to assess the reliability of the interactions between subunits. A pTM score above 0.8 indicates high-confidence and high-quality predictions, and scores below 0.6 typically reflect failed predictions. If the initial prediction displayed a well-structured core with disordered C– or N-terminal regions, the disordered segments were removed, and the trimmed sequence was re-submitted to AlphaFold 3. Various multimerization states were also tested with AlphaFold 3.

Traced models, or predictions with considerable reliability (pTM>0.75), were sent to DALI (PDB25 comparison) (Holm, 2022) and Foldseek (PDB100) servers (van Kempen et al., 2024) for structural homolog searches. Both results were considered to find a consensus. Even though it has been reported that DALI, although slower, it is more sensitive than Foldseek (Nomburg, Doherty et al., 2024).

Additional sequence-based software were used: InterPro for classification of protein families (Blum et al., 2025) and Contrastive Learning enabled Enzyme Annotation (CLEAN) to predict enzyme commission (EC) numbers (Yu et al., 2023).

## Data availability

The NO16 mass spectrometry data have been deposited to the ProteomeXchange Consortium (https://www.proteomexchange.org/) via the PRIDE partner repository (https://www.ebi.ac.uk/pride/) (Perez-Riverol, Bandla et al., 2025) with dataset identifier PXD070644.

The NO16 cryo-EM map and molecular models have been deposited at the Electron Microscopy Data Bank (EMDB, http://www.ebi.ac.uk/pdbe/emdb) and Protein Data Bank (PDB, https://www.ebi.ac.uk/pdbe/) with accession numbers EMD-55728 and 9T9R (vertex structure from localized reconstruction), EMD-55739 and 9T9V (icosahedral capsid structure). The validation reports are included as **Supplementary Files 4 and 5**.

## Supporting information

Supplementary file with supplementary figures and tables

## Acknowledgements

Work supported by grants from the Spanish State Research Agency (AEI/10.13039/501100011033), with co-funding from the European Regional Development Fund (PID2019-104098GB-I00 and PID2022-136456NB-I00) to C.S.M.. The CNB-CSIC was further supported by AEI Severo Ochoa Excellence grants SEV-2017-0712 and CEX2023-001386-S). S.O.U. held a CSIC JAE Severo Ochoa and Maria de Maeztu fellowship (JAE-SOMdM20-20) and a predoctoral contract from the Spanish Ministry of Science, Innovation and Universities (FPU2020-05148). M.Mi. was financially supported by grants from European Union under the Horizon Europe Programme, Grant Agreement No. 101084204 (Cure4Aqua), from the Innovation Fund Denmark project No. 2105-00014B (AQUAPHAGE) and the Danish National Research Foundation through the Danish Center for Hadal Research (HADAL, Grant No. DNRF145).

We thank Miguel Marcin from the CNB-CSIC Proteomics facility; Beatriz Martín, Javier Chichón and Noelia Zamarreño from the CNB-CSIC Electron and Cryo-electron Microscopy facilities for the excellent technical support; as well as Eilis Bragginton from Diamond Light Source for Titan Krios data collection at the UK national electron Bio-Imaging Center (eBIC) under proposal BI30374-14.

## Author contributions

Research design has been accomplished by S.O.U., G.N.C. and C.S.M., and experimental research has been completed mainly by S.O.U., helped by G.N.C. and P.K. in the early stages of the project. P.K. and M.Mi. contributed with relevant insight about the virus biology. Cryo-EM workflow for icosahedral averaged map has been processed by S.O.U., helped by G.N.C. and C.S.M.. M.Ma. obtained local reconstruction maps. S.O.U. traced and analysed the results. S.O.U., G.N.C., M.Ma. and C.S.M wrote the paper with contributions from all authors.

## Competing interests statement

The authors declare no competing interests.

## Notes

### Competing Interest Statement

The authors have declared no competing interest.

## References

1. Abdelrahman F, Easwaran M, Daramola OI, Ragab S, Lynch S, Oduselu TJ, Khan FM, Ayobami A, Adnan F, Torrents E, Sanmukh S, El-Shibiny A (2021) Phage-encoded endolysins. Antibiotics (Basel) 10

2. Abramson J, Adler J, Dunger J, Evans R, Green T, Pritzel A, Ronneberger O, Willmore L, Ballard AJ, Bambrick J, Bodenstein SW, Evans DA, Hung CC, O’Neill M, Reiman D, Tunyasuvunakool K, Wu Z, Zemgulyte A, Arvaniti E, Beattie C et al. (2024) Accurate structure prediction of biomolecular interactions with AlphaFold 3. Nature 630: 493–500

3. Abrescia NG, Cockburn JJ, Grimes JM, Sutton GC, Diprose JM, Butcher SJ, Fuller SD, San Martín C, Burnett RM, Stuart DI, Bamford DH, Bamford JK (2004) Insights into assembly from structural analysis of bacteriophage PRD1. Nature 432: 68–74

4. Abrescia NG, Grimes JM, Kivela HM, Assenberg R, Sutton GC, Butcher SJ, Bamford JK, Bamford DH, Stuart DI (2008) Insights into virus evolution and membrane biogenesis from the structure of the marine lipid-containing bacteriophage PM2. Mol Cell 31: 749–61

5. Abrishami V, Ilca SL, Gomez-Blanco J, Rissanen I, de la Rosa-Trevin JM, Reddy VS, Carazo JM, Huiskonen JT (2021) Localized reconstruction in Scipion expedites the analysis of symmetry mismatches in cryo-EM data. Prog Biophys Mol Biol 160: 43–52

6. Bae B, Ohene-Adjei S, Kocherginskaya S, Mackie RI, Spies MA, Cann IK, Nair SK (2008) Molecular basis for the selectivity and specificity of ligand recognition by the family 16 carbohydrate-binding modules from Thermoanaerobacterium polysaccharolyticum ManA. J Biol Chem 283: 12415–25

7. Bannister AJ, Kouzarides T (2011) Regulation of chromatin by histone modifications. Cell Res 21: 381–95

8. Bardy P, MacDonald CIW, Pantucek R, Antson AA, Fogg PCM (2023) Jorvik: A membrane-containing phage that will likely found a new family within Vinavirales. iScience 26: 108104

9. Bewley MC, Springer K, Zhang Y-B, Freimuth P, Flanagan JM (1999) Structural analysis of the mechanism of adenovirus binding to its human cellular receptor, CAR. Science 286: 1579–83

10. Blum M, Andreeva A, Florentino LC, Chuguransky SR, Grego T, Hobbs E, Pinto BL, Orr A, Paysan-Lafosse T, Ponamareva I, Salazar GA, Bordin N, Bork P, Bridge A, Colwell L, Gough J, Haft DH, Letunic I, Llinares-Lopez F, Marchler-Bauer A et al. (2025) InterPro: the protein sequence classification resource in 2025. Nucleic Acids Res 53: D444–D456

11. Brewer GJ (1978) Membrane-localized replication of bacteriophage PM2. Virology 84: 242–5

12. Brum JR, Schenck RO, Sullivan MB (2013) Global morphological analysis of marine viruses shows minimal regional variation and dominance of non-tailed viruses. ISME J 7: 1738–51

13. Byrne MJ, Iadanza MG, Perez MA, Maskell DP, George RM, Hesketh EL, Beales PA, Zack MD, Berry C, Thompson RF (2021) Cryo-EM structures of an insecticidal Bt toxin reveal its mechanism of action on the membrane. Nat Commun 12: 2791

14. Caspar DL, Klug A (1962) Physical principles in the construction of regular viruses. Cold Spring Harb Symp Quant Biol 27: 1–24

15. Cernooka E, Rumnieks J, Zrelovs N, Tars K, Kazaks A (2022) Diversity of the lysozyme fold: structure of the catalytic domain from an unusual endolysin encoded by phage Enc34. Sci Rep 12: 5005

16. Chiu CY, Wu E, Brown SL, Von Seggern DJ, Nemerow GR, Stewart PL (2001) Structural analysis of a fiber-pseudotyped adenovirus with ocular tropism suggests differential modes of cell receptor interactions. J Virol 75: 5375–80

17. Christensen DG, Xie X, Basisty N, Byrnes J, McSweeney S, Schilling B, Wolfe AJ (2019) Post-translational protein acetylation: an elegant mechanism for bacteria to dynamically regulate metabolic functions. Front Microbiol 10: 1604

18. Ciordia S, Alvarez-Sola G, Rullan M, Urman JM, Avila MA, Corrales FJ (2021) Digging deeper into bile proteome. J Proteomics 230: 103984

19. Condezo GN, San Martín C (2017) Localization of adenovirus morphogenesis players, together with visualization of assembly intermediates and failed products, favor a model where assembly and packaging occur concurrently at the periphery of the replication center. PLoS Pathog 13: e1006320

20. Cuervo A, Dauden MI, Carrascosa JL (2013) Nucleic acid packaging in viruses. Sub-cellular biochemistry 68: 361–94

21. Cvirkaite-Krupovic V, Krupovic M, Daugelavicius R, Bamford DH (2010) Calcium ion-dependent entry of the membrane-containing bacteriophage PM2 into its Pseudoalteromonas host. Virology 405: 120–8

22. de la Rosa-Trevin JM, Oton J, Marabini R, Zaldivar A, Vargas J, Carazo JM, Sorzano CO (2013) Xmipp 3.0: an improved software suite for image processing in electron microscopy. J Struct Biol 184: 321–8

23. de la Rosa-Trevin JM, Quintana A, Del Cano L, Zaldivar A, Foche I, Gutierrez J, Gomez-Blanco J, Burguet-Castell J, Cuenca-Alba J, Abrishami V, Vargas J, Oton J, Sharov G, Vilas JL, Navas J, Conesa P, Kazemi M, Marabini R, Sorzano CO, Carazo JM (2016) Scipion: A software framework toward integration, reproducibility and validation in 3D electron microscopy. J Struct Biol 195: 93–9

24. Emsley P, Lohkamp B, Scott WG, Cowtan K (2010) Features and development of Coot. Acta Crystallogr D Biol Crystallogr 66: 486–501

25. Espejo RT, Canelo ES, Sinsheimer RL (1971) Replication of bacteriophage PM2 deoxyribonucleic acid: a closed circular double-stranded molecule. J Mol Biol 56: 597–621

26. Favrot L, Blanchard JS, Vergnolle O (2016) Bacterial GCN5-related N-acetyltransferases: from resistance to regulation. Biochemistry 55: 989–1002

27. Frans I, Michiels CW, Bossier P, Willems KA, Lievens B, Rediers H (2011) Vibrio anguillarum as a fish pathogen: virulence factors, diagnosis and prevention. J Fish Dis 34: 643–61

28. Gaidelyte A, Cvirkaite-Krupovic V, Daugelavicius R, Bamford JK, Bamford DH (2006) The entry mechanism of membrane-containing phage Bam35 infecting Bacillus thuringiensis. J Bacteriol 188: 5925–34

29. Garber EA, Seidman MM, Levine AJ (1980) Intracellular SV40 nucleoprotein complexes: synthesis to encapsidation. Virology 107: 389–401

30. Gómez-Gónzalez A, Burkhardt P, Bauer M, Suomalainen M, Mateos JM, Loehr MO, Luedtke NW, Greber UF (2024) Stepwise virus assembly in the cell nucleus revealed by spatiotemporal click chemistry of DNA replication. Sci Adv 10: eadq7483

31. Happonen LJ, Oksanen E, Liljeroos L, Goldman A, Kajander T, Butcher SJ (2013) The structure of the NTPase that powers DNA packaging into Sulfolobus turreted icosahedral virus 2. J Virol 87: 8388–98

32. Hartman R, Eilers BJ, Bollschweiler D, Munson-McGee JH, Engelhardt H, Young MJ, Lawrence CM (2019) The molecular mechanism of cellular attachment for an archaeal virus. Structure 27: 1634–1646 e3

33. Hocher A, Laursen SP, Radford P, Tyson J, Lambert C, Stevens KM, Montoya A, Shliaha PV, Picardeau M, Sockett RE, Luger K, Warnecke T (2023) Histones with an unconventional DNA-binding mode in vitro are major chromatin constituents in the bacterium Bdellovibrio bacteriovorus. Nat Microbiol 8: 2006–2019

34. Holm L (2022) Dali server: structural unification of protein families. Nucleic Acids Res 50: W210–W215

35. Hong C, Oksanen HM, Liu X, Jakana J, Bamford DH, Chiu W (2014) A structural model of the genome packaging process in a membrane-containing double stranded DNA virus. PLoS biology 12: e1002024

36. Huiskonen JT, Kivela HM, Bamford DH, Butcher SJ (2004) The PM2 virion has a novel organization with an internal membrane and pentameric receptor binding spikes. Nat Struct Mol Biol 11: 850–6

37. Huiskonen JT, Manole V, Butcher SJ (2007) Tale of two spikes in bacteriophage PRD1. Proc Natl Acad Sci U S A 104: 6666–71

38. Jamali K, Kall L, Zhang R, Brown A, Kimanius D, Scheres SHW (2024) Automated model building and protein identification in cryo-EM maps. Nature 628: 450–457

39. Jumper J, Evans R, Pritzel A, Green T, Figurnov M, Ronneberger O, Tunyasuvunakool K, Bates R, Zidek A, Potapenko A, Bridgland A, Meyer C, Kohl SAA, Ballard AJ, Cowie A, Romera-Paredes B, Nikolov S, Jain R, Adler J, Back T et al. (2021) Highly accurate protein structure prediction with AlphaFold. Nature 596: 583–589

40. Kalatzis PG, Carstens AB, Katharios P, Castillo D, Hansen LH, Middelboe M (2019) Complete Genome Sequence of Vibrio anguillarum Nontailed Bacteriophage NO16. Microbiol Resour Announc 8

41. Kalatzis PG, Mauritzen JJ, Winther-Have CS, Michniewski S, Millard A, Tsertou MI, Katharios P, Middelboe M (2023) Staying below the radar: unraveling a new family of ubiquitous “cryptic” non-tailed temperate vibriophages and implications for their bacterial hosts. Int J Mol Sci 24

42. Kauffman KM, Hussain FA, Yang J, Arevalo P, Brown JM, Chang WK, VanInsberghe D, Elsherbini J, Sharma RS, Cutler MB, Kelly L, Polz MF (2018) A major lineage of non-tailed dsDNA viruses as unrecognized killers of marine bacteria. Nature 554: 118–122

43. Kejzar N, Laanto E, Rissanen I, Abrishami V, Selvaraj M, Moineau S, Ravantti J, Sundberg LR, Huiskonen JT (2022) Cryo-EM structure of ssDNA bacteriophage PhiCjT23 provides insight into early virus evolution. Nat Commun 13: 7478

44. Kivela HM, Kalkkinen N, Bamford DH (2002) Bacteriophage PM2 has a protein capsid surrounding a spherical proteinaceous lipid core. J Virol 76: 8169–78

45. Kivela HM, Mannisto RH, Kalkkinen N, Bamford DH (1999) Purification and protein composition of PM2, the first lipid-containing bacterial virus to be isolated. Virology 262: 364–74

46. Koenig T, Menze BH, Kirchner M, Monigatti F, Parker KC, Patterson T, Steen JJ, Hamprecht FA, Steen H (2008) Robust prediction of the MASCOT score for an improved quality assessment in mass spectrometric proteomics. J Proteome Res 7: 3708–17

47. Kong AT, Leprevost FV, Avtonomov DM, Mellacheruvu D, Nesvizhskii AI (2017) MSFragger: ultrafast and comprehensive peptide identification in mass spectrometry-based proteomics. Nat Methods 14: 513–520

48. Koonin EV, Dolja VV, Krupovic M (2022) The logic of virus evolution. Cell Host Microbe 30: 917–929

49. Koonin EV, Dolja VV, Krupovic M, Varsani A, Wolf YI, Yutin N, Zerbini FM, Kuhn JH (2020) Global Organization and Proposed Megataxonomy of the Virus World. Microbiol Mol Biol Rev 84

50. Krissinel E, Henrick K (2007) Inference of macromolecular assemblies from crystalline state. J Mol Biol 372: 774–97

51. Krogh A, Larsson B, von Heijne G, Sonnhammer EL (2001) Predicting transmembrane protein topology with a hidden Markov model: application to complete genomes. J Mol Biol 305: 567–80

52. Krupovic M, Bamford DH (2007) Putative prophages related to lytic tailless marine dsDNA phage PM2 are widespread in the genomes of aquatic bacteria. BMC Genomics 8: 236

53. Krupovic M, Koonin EV (2017) Multiple origins of viral capsid proteins from cellular ancestors. Proc Natl Acad Sci U S A 114: E2401–E2410

54. Krupovic M, Makarova KS, Koonin EV (2022) Cellular homologs of the double jelly-roll major capsid proteins clarify the origins of an ancient virus kingdom. Proc Natl Acad Sci U S A 119

55. Kukkaro P, Bamford DH (2009) Virus-host interactions in environments with a wide range of ionic strengths. Environ Microbiol Rep 1: 71–7

56. Laanto E, Mantynen S, De Colibus L, Marjakangas J, Gillum A, Stuart DI, Ravantti JJ, Huiskonen JT, Sundberg LR (2017) Virus found in a boreal lake links ssDNA and dsDNA viruses. Proc Natl Acad Sci U S A 114: 8378–8383

57. Lasso G, Honig B, Shapira SD (2021) A Sweep of Earth’s Virome Reveals Host-Guided Viral Protein Structural Mimicry and Points to Determinants of Human Disease. Cell Syst 12: 82–91 e3

58. Laurinmaki PA, Huiskonen JT, Bamford DH, Butcher SJ (2005) Membrane proteins modulate the bilayer curvature in the bacterial virus Bam35. Structure 13: 1819–28

59. Lee SY, Kim GE, Park HH (2022) Molecular basis of transcriptional repression of anti-CRISPR by anti-CRISPR-associated 2. Acta Crystallogr D Struct Biol 78: 59–68

60. Liebschner D, Afonine PV, Baker ML, Bunkoczi G, Chen VB, Croll TI, Hintze B, Hung LW, Jain S, McCoy AJ, Moriarty NW, Oeffner RD, Poon BK, Prisant MG, Read RJ, Richardson JS, Richardson DC, Sammito MD, Sobolev OV, Stockwell DH et al. (2019) Macromolecular structure determination using X-rays, neutrons and electrons: recent developments in Phenix. Acta Crystallogr D Struct Biol 75: 861–877

61. Liu Y, Zhang L, Guo M, Chen L, Wu B, Huang H (2021) Structural basis for anti-CRISPR repression mediated by bacterial operon proteins Aca1 and Aca2. J Biol Chem 297: 101357

62. Madeira F, Madhusoodanan N, Lee J, Eusebi A, Niewielska A, Tivey ARN, Lopez R, Butcher S (2024) The EMBL-EBI Job Dispatcher sequence analysis tools framework in 2024. Nucleic Acids Res 52: W521–W525

63. Mannisto RH, Kivela HM, Paulin L, Bamford DH, Bamford JK (1999) The complete genome sequence of PM2, the first lipid-containing bacterial virus to be isolated. Virology 262: 355–63

64. Mantynen S, Laanto E, Sundberg LR, Poranen MM, Oksanen HM, Report Consortium I (2020) ICTV Virus Taxonomy Profile: Finnlakeviridae. J Gen Virol 101: 894–895

65. Martínez M, Gallardo J, Condezo GN, San Martín C (2025) Chapter 1 – Structural biology of adenovirus. In Adenoviral Vectors for Gene Therapy (Third Edition), Curiel DT, Parker AL (eds) pp 1–43. San Diego: Academic Press

66. Martínez M, Jiménez-Moreno A, Maluenda D, Ramírez-Aportela E, Melero R, Cuervo A, Conesa P, del Caño L, Fonseca YC, Sánchez-García R, Strelak D, Conesa JJ, Fernández-Giménez E, de Isidro F, Sorzano COS, Carazo JM, Marabini R (2020) Integration of cryo-EM model building software in Scipion. Journal of Chemical Information and Modeling 60: 2533–2540

67. Meng EC, Goddard TD, Pettersen EF, Couch GS, Pearson ZJ, Morris JH, Ferrin TE (2023) UCSF ChimeraX: Tools for structure building and analysis. Protein Sci 32: e4792

68. Merckel MC, Huiskonen JT, Bamford DH, Goldman A, Tuma R (2005) The structure of the bacteriophage PRD1 spike sheds light on the evolution of viral capsid architecture. Mol Cell 18: 161–70

69. Nomburg J, Doherty EE, Price N, Bellieny-Rabelo D, Zhu YK, Doudna JA (2024) Birth of protein folds and functions in the virome. Nature 633: 710–717

70. Núñez-Ramírez R, Huesa J, Bel Y, Ferre J, Casino P, Arias-Palomo E (2020) Molecular architecture and activation of the insecticidal protein Vip3Aa from Bacillus thuringiensis. Nat Commun 11: 3974

71. Orsburn BC (2021) Proteome Discoverer-a community enhanced data processing suite for protein informatics. Proteomes 9

72. Overton MS, Manuel RD, Lawrence CM, Snyder JC (2023) Viruses of the Turriviridae: an emerging model system for studying archaeal virus-host interactions. Front Microbiol 14: 1258997

73. Peralta B, Gil-Carton D, Castano-Diez D, Bertin A, Boulogne C, Oksanen HM, Bamford DH, Abrescia NG (2013) Mechanism of membranous tunnelling nanotube formation in viral genome delivery. PLoS biology 11: e1001667

74. Pérez-Berná AJ, Mangel WF, McGrath WJ, Graziano V, Flint J, San Martín C (2014) Processing of the L1 52/55k protein by the adenovirus protease: a new substrate and new insights into virion maturation. J Virol 88: 1513–24

75. Perez-Riverol Y, Bandla C, Kundu DJ, Kamatchinathan S, Bai J, Hewapathirana S, John NS, Prakash A, Walzer M, Wang S, Vizcaino JA (2025) The PRIDE database at 20 years: 2025 update. Nucleic Acids Res 53: D543–D553

76. Pettersen EF, Goddard TD, Huang CC, Meng EC, Couch GS, Croll TI, Morris JH, Ferrin TE (2021) UCSF ChimeraX: Structure visualization for researchers, educators, and developers. Protein Sci 30: 70–82

77. Prevelige PE, Fane BA (2012) Building the machines: scaffolding protein functions during bacteriophage morphogenesis. Advances in experimental medicine and biology 726: 325–50

78. Punjani A, Rubinstein JL, Fleet DJ, Brubaker MA (2017) cryoSPARC: algorithms for rapid unsupervised cryo-EM structure determination. Nat Methods 14: 290–296

79. Reddy HK, Carroni M, Hajdu J, Svenda M (2019) Electron cryo-microscopy of bacteriophage PR772 reveals the elusive vertex complex and the capsid architecture. eLife 8

80. Robert X, Gouet P (2014) Deciphering key features in protein structures with the new ENDscript server. Nucleic Acids Res 42: W320–4

81. Rohou A, Grigorieff N (2015) CTFFIND4: Fast and accurate defocus estimation from electron micrographs. J Struct Biol 192: 216–21

82. Russo CJ, Henderson R (2018) Ewald sphere correction using a single side-band image processing algorithm. Ultramicroscopy 187: 26–33

83. Rydman PS, Bamford DH (2000) Bacteriophage PRD1 DNA entry uses a viral membrane-associated transglycosylase activity. Mol Microbiol 37: 356–63.

84. San Martín C, van Raaij MJ (2018) The so far farthest reaches of the double jelly roll capsid protein fold. Virol J 15: 181

85. Scheres SH (2012) RELION: implementation of a Bayesian approach to cryo-EM structure determination. J Struct Biol 180: 519–30

86. Shevchenko A, Wilm M, Vorm O, Mann M (1996) Mass spectrometric sequencing of proteins silver-stained polyacrylamide gels. Analytical chemistry 68: 850–8

87. Snijder J, Borst AJ, Dosey A, Walls AC, Burrell A, Reddy VS, Kollman JM, Veesler D (2017) Vitrification after multiple rounds of sample application and blotting improves particle density on cryo-electron microscopy grids. J Struct Biol 198: 38–42

88. Stayrook S, Jaru-Ampornpan P, Ni J, Hochschild A, Lewis M (2008) Crystal structure of the lambda repressor and a model for pairwise cooperative operator binding. Nature 452: 1022–5

89. Steensen K, Seneca J, Bartlau N, Yu XA, Hussain FA, Polz MF (2024) Tailless and filamentous prophages are predominant in marine Vibrio. ISME J 18

90. Turner D, Adriaenssens EM, Amann RI, Bardy P, Bartlau N, Barylski J, Blazejak S, Bouzari M, Briegel A, Briers Y, Carrillo D, Chen X, Claessen D, Cook R, Crisci MA, Dechesne A, Deptula P, Dutilh BE, Ely B, Fieseler L et al. (2025) Summary of taxonomy changes ratified by the International Committee on Taxonomy of Viruses (ICTV) from the Bacterial Viruses Subcommittee, 2025. J Gen Virol 106

91. van Kempen M, Kim SS, Tumescheit C, Mirdita M, Lee J, Gilchrist CLM, Soding J, Steinegger M (2024) Fast and accurate protein structure search with Foldseek. Nat Biotechnol 42: 243–246

92. Vilas JL, Gomez-Blanco J, Conesa P, Melero R, Miguel de la Rosa-Trevin J, Oton J, Cuenca J, Marabini R, Carazo JM, Vargas J, Sorzano COS (2018) MonoRes: Automatic and Accurate Estimation of Local Resolution for Electron Microscopy Maps. Structure 26: 337–344 e4

93. Wu E, Pache L, Von Seggern DJ, Mullen TM, Mikyas Y, Stewart PL, Nemerow GR (2003) Flexibility of the adenovirus fiber is required for efficient receptor interaction. J Virol 77: 7225–35

94. Xu L, Benson SD, Butcher SJ, Bamford DH, Burnett RM (2003) The receptor binding protein P2 of PRD1, a virus targeting antibiotic-resistant bacteria, has a novel fold suggesting multiple functions. Structure 11: 309–322

95. Yu T, Cui H, Li JC, Luo Y, Jiang G, Zhao H (2023) Enzyme function prediction using contrastive learning. Science 379: 1358–1363

96. Yutin N, Backstrom D, Ettema TJG, Krupovic M, Koonin EV (2018) Vast diversity of prokaryotic virus genomes encoding double jelly-roll major capsid proteins uncovered by genomic and metagenomic sequence analysis. Virol J 15: 67

97. Yutin N, Rayko M, Antipov D, Mutz P, Wolf YI, Krupovic M, Koonin EV (2022) Varidnaviruses in the human gut: a major expansion of the order Vinavirales. Viruses 14

98. Zheng SQ, Palovcak E, Armache JP, Verba KA, Cheng Y, Agard DA (2017) MotionCor2: anisotropic correction of beam-induced motion for improved cryo-electron microscopy. Nat Methods 14: 331–332

99. Zivanov J, Nakane T, Forsberg BO, Kimanius D, Hagen WJ, Lindahl E, Scheres SH (2018) New tools for automated high-resolution cryo-EM structure determination in RELION-3. Elife 7

