## Supplementary file with supplementary figures and tables for "Structure of NO16, a marine non-tailed vibriophage with a global distribution"

**Short title: Structure of bacteriophage NO16**

**The PDF file includes:**

**Supplementary tables S1 to S6**

**Supplementary figures S1 to S10**

**Supplementary references**

**Additional supplementary files and movie, not included in this PDF:**

**Supplementary File 1 (.xlsx).** Mass spectrometry results of NO16: (a) 60 kDa contaminant band. (b) purified NO16 virion.

**Supplementary File 2 (.xlsx).** Structural homology search results of NO16 proteins.

**Supplementary File 3 (.json).** Local reconstruction workflow.

**Supplementary File 4 (.pdf).** Validation report for NO16 icosahedral capsid structure.

**Supplementary File 5 (.pdf).** Validation report for NO16 localized reconstruction vertex structure.

**Supplementary Movie 1.** Summary of NO16 structure.

### Supplementary Tables

**Table S1. Cryo-EM vitrification conditions, data collection, image processing, reconstruction and refinement.**

| **Vitrification** | | |
|---|---|---|
| Grids | Quantifoil R1.2/1.3 Cu/Rh | |
| Glow discharge | 25 mA, 1 minute | |
| Temperature | 22ºC | |
| Humidity | 95% | |
| Blot time | 3 seconds | |
| Force | - 4 | |
| **Data collection** | | |
| Microscope | Titan Krios | |
| Camera | Bioquantum K3 | |
| Voltage | 300 kV | |
| Magnification | 64,000 x | |
| Nominal pixel size | 1.34 Å/px | |
| Exposure time | 4.66 s | |
| Total dose rate | 39.97 e-/Å$^2$ | |
| Number of frames | 40 | |
| Defocus range | -1.1 to -2.5 µm, in -0.2 µm steps | |
| Micrographs collected | 20,640 | |
| Acquisition software | EPU 3.5.1 | |
| **Image processing** | | |
| Frame alignment software | MotionCor2, Scipion 3 | |
| CTF estimation software | CTFFIND4, Scipion 3 | |
| Particle picking software | Xmipp3, Scipion 3 | |
| Micrographs used | 20,526 | |
| Particles picked | 18,657 | |
| **Reconstruction** | **Whole capsid** | **Local reconstruction** |
| Software | RELION, Scipion 3 | RELION, CryoSPARC |
| Particles included | 15,429 | 162,312 (sub-particles) |
| Symmetry imposed | Icosahedral | C1 |
| Rotational accuracy | 0.029 degrees | 1.89 degrees |
| Translational accuracy | 0.1005 pixels | 1.24 pixels |
| B-factor applied | -108.24 Å$^2$ | -131.28 Å$^2$ |
| Final resolution (gold standard FSC = 0.143) | 3.3 Å | 3.9 Å |

**Table S2. Modelling and validation statistics for NO16 molecular models and maps.** Parameters provided by Phenix real-space refine (Liebschner, Afonine et al., 2019).

| | **Icosahedral map** | **Local reconstructed map** |
|---|---|---|
| Software | Scipion, ModelAngelo, ChimeraX, Coot, Phenix | |
| Model | PDB ID 9T9V | PDB ID 9T9R |
| Chains | 19 | 8 |
| Atoms | 22065 (Hydrogens: 0) | 6877 (Hydrogens: 0) |
| Residues | Protein: 2864 Nucleotide: 0 | Protein: 893 Nucleotide: 0 |
| Water | 0 | 0 |
| Ligands | 4 | 0 |
| Bond length (RMSD, Å) | 0.004 | 0.007 |
| Bond angles (RMSD, °) | 0.97 | 1.205 |
| MolProbity score | 1.73 | 2.35 |
| Clash score | 7.62 | 17.12 |
| **Ramachandran plot (%)** | | |
| Outliers | 0.04 | 0.36 |
| Allowed | 4.41 | 12.46 |
| Favoured | 95.56 | 87.19 |
| **Rama-Z (Ramachandran plot Z-score, RMSD)** | | |
| whole | (N= 2834) -0.98 (0.16) | (N= 835) -2.96 (0.27) |
| helix | (N= 405) 1.13 (0.28) | (N= 18) -3.03 (1.09) |
| sheet | (N= 998) -0.58 (0.17) | (N= 179) -0.98 (0.43) |
| loop | (N= 1432) -1.19 (0.16) | (N= 638) -2.44 (0.22) |
| Rotamer outliers (%) | 0.17 | 0.77 |
| Cβ outliers (%) | 0 | 0 |

**Table S3. NO16 proteins traced in the model.** In grey, chains traced in the local reconstructed map.

| Protein | Length (amino acids) | Copy number in the AU* | Chain ID | Residues traced | Not traced |
|---|---|---|---|---|---|
| **Icosahedral AU** | | | | | |
| **MCP (GP19)** | 265 | 10 | A | 2-265 | 1 |
| | | | B | 2-265 | 1 |
| | | | C | 2-265 | 1 |
| | | | D | 2-265 | 1 |
| | | | E | 2-265 | 1 |
| | | | F | 2-265 | 1 |
| | | | G | 2-265 | 1 |
| | | | H | 2-265 | 1 |
| | | | I | 2-265 | 1 |
| | | | J | 2-265 | 1 |
| | Cation | 4 | P | UNX | - |
| | | | Q | UNX | - |
| | | | R | UNX | - |
| | | | S | UNX | - |
| **Penton base proximal domain (GP14) 1-88 aa** | 180 | 1 | K | 2-88 | Distal domain traced in PDB vertex |
| **Minor capsid protein (GP18)** | 82 | 3 | L | 4-50 | 1-3; 51-82 |
| | | | M | 9-42 | 1-8; 43-82 |
| | | | N | 2-55 | 1; 56-82 |
| **Unknown** | - | 1 | O | 1-12 (UNK) | - |
| **Non-icosahedral vertex** | | | | | |
| **Penton base distal domain (GP14) 89-180aa** | 180 | 1 | A | 66-69, 89-180 | Proximal domain in PDB AU |
| | | | B | 89-131,143-180 | 132-142 |
| | | | C | 89-99,104-131, 143-149, 159-180 | 100-103,132-142,150-158 |
| | | | D | 89-101, 108-130,141-163,172-180 | 102-107,131-140,164-171 |
| | | | E | 89-101,110-129,139-152, 171-180 | 102-109,130-138,153-170 |
| **Spike (GP13)** | 235 | SM** | F | 11-235 | 1-10 |
| | | SM | G | 8-36,45-60,81-88, 137-225, 230-235 | 1-8,37-44,61-80,89-136,226-229 |
| | | SM | H | 7-36, 45-59, 80-87, 136-225, 230-235 | 1-6,37-44,60-79,88-135,226-229 |

*AU: icosahedral asymmetric unit. **SM: Symmetry mismatch.

**Table S4. PDBePISA (Krissinel & Henrick, 2007) analysis of NO16 capsid proteins.** Chains colored by position in the asymmetric unit, following color legend of Fig. 1c. Interface area (Å$^2$) is calculated as half the difference between the total accessible surface areas of the isolated and interacting structures. ΔiG represents the solvation-free energy gain upon interface formation, it is calculated as the difference in total solvation energies between the isolated and interacting structures, and a negative ΔiG indicates favorable protein-protein interaction. $N_{HB}$ indicates the number of potential hydrogen bonds formed across the interface. $N_{SB}$ indicates the number of potential salt bridges across the interface. $N_{DS}$ indicates the number of disulfide bounds across the interface. Notice that PDBePISA considers interactions between atoms, so some of the salt bridge atoms could correspond to the same residue.

| Chain 1 (capsomer position) | Chain 2 (capsomer position) | Interface area (Å$^2$) | ΔiG (kcal/mol) | $N_{HB}$ | $N_{SB}$ | $N_{DS}$ |
|---|---|---|---|---|---|---|
| Intra-capsomer contacts between MCP | | | | | | |
| J (4) | H (4) | 852.4 | -2.3 | 4 | 11 | 0 |
| F (2) | D (2) | 836.9 | -1.0 | 5 | 12 | 0 |
| F (2) | E (2) | 836.4 | -4.4 | 4 | 9 | 0 |
| E (2) | D (2) | 818.1 | -4.4 | 6 | 10 | 0 |
| J (4) | I (4) | 816.3 | -3.9 | 8 | 11 | 0 |
| I (4) | H (4) | 815.8 | -2.9 | 5 | 9 | 0 |
| B (1) | A (1) | 796.0 | -3.5 | 2 | 6 | 0 |
| C (1) | B (1) | 783.8 | -2.0 | 6 | 8 | 0 |
| C (1) | A (1) | 636.7 | 2.2 | 2 | 8 | 0 |
| Inter-capsomer contacts between MCP | | | | | | |
| H (4) | C (1) | 584.9 | -1.0 | 5 | 1 | 0 |
| I (4) | F (2) | 425.9 | -1.2 | 1 | 1 | 0 |
| I (4) | G (3) | 395.5 | -2.0 | 0 | 0 | 0 |
| D (2) | B (1) | 389.4 | -4.0 | 0 | 0 | 0 |
| G (3) | F (2) | 378.3 | -0.4 | 0 | 2 | 0 |
| I (4) | C (1) | 230.0 | 0.7 | 1 | 0 | 0 |
| I (4) | D (2) | 159.4 | 1.1 | 1 | 0 | 0 |
| D (2) | C (1) | 156.9 | 1.2 | 0 | 0 | 0 |
| J (4) | G (3) | 137.6 | 0.3 | 2 | 0 | 0 |
| Contacts between MCP-penton | | | | | | |
| K | A (1) | 719.2 | -2.5 | 2 | 5 | 0 |
| Contacts between spike-penton | | | | | | |
| F (GP13)* | A (GP14)* | 590.7 | -6.0 | 4 | 0 | 0 |
| Contacts between MCP-mCP | | | | | | |
| N | J (4) | 1408.8 | -19.3 | 7 | 2 | 0 |
| L | C (1) | 1311.7 | -14.8 | 6 | 2 | 0 |
| M | D (2) | 961.1 | -15.2 | 2 | 1 | 0 |
| M | F (2) | 346.6 | -0.6 | 2 | 6 | 0 |
| N | I (4) | 327.4 | -0.9 | 4 | 4 | 0 |
| L | B (1) | 317.7 | -1.7 | 0 | 3 | 0 |
| N | G (3) | 314.0 | 0.7 | 2 | 0 | 0 |
| L | D (2) | 173.2 | -2.0 | 0 | 0 | 0 |
| N | H (4) | 210.9 | -4.7 | 1 | 0 | 0 |
| L | A (1) | 196.6 | -2.9 | 1 | 0 | 0 |
| O | H (4) | 197.5 | -3.7 | 1 | 0 | 0 |
| L | I (4) | 197.2 | -0.2 | 5 | 0 | 0 |
| O | I (4) | 149.0 | -2.9 | 0 | 0 | 0 |
| M | I (4) | 95.2 | -0.7 | 0 | 0 | 0 |
| O | C (1) | 26.3 | -0.5 | 0 | 0 | 0 |

*From the non-icosahedral vertex molecular model.

**Table S5. PDBePISA analysis of PM2, FLIP and ØCJT23 MCP capsomers.** For comparison with Table S4. Parameters are descripted in Table S4.

| | Chain 1 | Chain 2 | Interface area (Å$^2$) | $\Delta^i G$ (kcal/mol) | $N_{HB}$ | $N_{SB}$ | $N_{DS}$ |
|---|---|---|---|---|---|---|---|
| **PM2** | A | B | 1164.9 | -6.7 | 21 | 10 | 0 |
| | B | C | 1159.1 | -7.7 | 21 | 10 | 0 |
| | A | C | 1153.8 | -7.5 | 21 | 10 | 0 |

| | Chain 1 | Chain 2 | Interface area (Å$^2$) | $\Delta^i G$ (kcal /mol) | $N_{HB}$ | $N_{SB}$ | $N_{DS}$ |
|---|---|---|---|---|---|---|---|
| **FLIP** | A | B | 1423.4 | -8.5 | 6 | 3 | 0 |
| | B | C | 1381.8 | -8.9 | 6 | 3 | 0 |
| | A | C | 1381.5 | -8.9 | 10 | 5 | 0 |

| | Chain 1 | Chain 2 | Interface area (Å$^2$) | $\Delta^i G$ (kcal /mol) | $N_{HB}$ | $N_{SB}$ | $N_{DS}$ |
|---|---|---|---|---|---|---|---|
| **ØCJT23** | A | B | 913.7 | -8.2 | 8 | 5 | 0 |
| | B | C | 913.5 | -8.9 | 10 | 6 | 0 |
| | A | C | 883.1 | -9.4 | 10 | 5 | 0 |

**Table S6. Structure prediction and homology search results for all proteins coded by NO16 genome.** MS, mass spectrometry. P, present in the virion in mass spectrometry analysis. pTM, predicted template modelling score for AlphaFold predictions. T, traced protein. IDR, intrinsically disordered region. TM, transmembrane. EC, Enzyme Commission number. Z-scores for homology using DALI range from 8 to 20, which indicate probable homology; values above 20 indicate definite homology; scores below 2 are considered nonsignificant. For Foldseek, E-value equal or lower than 0.01 was considered. Proteins for which a putative function has been assigned are highlighted in green.

| **Protein** | **Previous annotation** (if not specified, unknown) | **MS** | **New annotation** | **pTM** | **Structural based search for homologues** | | **Sequence based software** | |
|---|---|---|---|---|---|---|---|---|
| | | | | | **DALI (PDB25)** | **Foldseek (PDB100)** | **InterPro** | **CLEAN** |
| **GP01** | | | | 0.72 | - | - | n/a | Fumarate hydratase EC:4.2.1.2 (low) |
| **GP02** | | | | 0.8 | <2 | >0.01 | n/a | Methyltransferase EC:2.1.1.86 (low) |
| **GP03** | | | Replication initiation protein | 0.78 | SIRV1 hypothetical protein ORF119, Replication initiation protein (PDB: 2x3g, Z-score: 9.3) | >0.01 | 340-409 aa Replication-associated protein ORF2/G2P (IPR056906) | DNA ligase (ATP) EC:6.5.1.1 (low) |
| **GP04** | | | | 0.26 | - | - | n/a | Deoxyribonuclease EC:3.1.21.4 (low) |
| **GP05** | | | | 0.39 | - | - | n/a | Decarboxylase EC:4.1.1.44 (low) |
| **GP06** | Putative DNA-binding protein | | Lytic and lysogenic transcriptional regulator | 0.84 | HTH-type transcription regulators (PDB: 2eby, Z-score: 7.7) | ImmR transcriptional regulator DNA-binding domain of *Bacillus subtilis* (PDB: 7t8i, E-value:5.17e-2) | n/a | Transferring alkyl or aryl groups EC:2.5.1.83 (low) |
| **GP07** | S-adenosylhomocysteine protein | | Anti-CRISPR-associated protein | 0.49 | Anti-CRISPR-associated Aca2 (PDB: 7ezy, Z-score: 10) | Anti-CRISPR-associated protein Aca2 (PDB: 7vjq, E-value: 1.80e-1) | 55-111 aa Phage protein (IPR021077) | Methyltransferases EC:2.1.1.148 (low) |
| **GP08** | | P | | 0.49 | - | - | 22-41 aa TM | Methyltransferases EC:2.1.1.86 (low) |
| **GP09** | | | | 0.38 | - | - | | Methyltransferases EC:2.1.1.86 (low) |
| **GP10** | Structurally similar to PM2 protein p6 | P | | 0.47 | - | - | | Thiosulfate dehydrogenase EC:1.8.5.2 (low) |
| **GP11** | Protein inside capsid D | P | Endolysin | 0.93 | Endolysin from bacteriophage Enc34 (PDB: 7q47, Z-score: 17.4) | Endolysin from bacteriophage Enc34 (PDB: 7q47, E-value: 1.712e-09) | 1-20 aa IDR | Lysozyme EC:3.2.1.17 (low) |

| Protein | **Previous annotation** (if not specified, unknown) | MS | New annotation | pTM | Structural based search for homologues | | Sequence based software | |
|---|---|---|---|---|---|---|---|---|
| | | | | | **DALI (PDB25)** | **Foldseek (PDB100)** | **InterPro** | **CLEAN** |
| **GP12** | | | | 0.64 | - | - | 40-64 aa IDR | Peptidase EC:3.4.22.65 (low) |
| **GP13** | | P | Peptidoglycan hydrolase | T | S-layer associated multidomain endoglucanase (PDB: 2zew, Z-score: 9.6) | Vip3Bc1 and Vip3Aa tetramer (PDB: 6yrf/6tfk, E-value: 2.29e-02, 4.63e-02) | n/a | Glycosylases EC:3.2.2.22 (low) |
| **GP14** | | P | Penton base protein | T | | | n/a | Carbon-carbon bonds ligase EC:6.4.1.8 (low) |
| **GP15** | | | | 0.36 | - | - | 128-145 aa TM | Methyltransferases EC:2.1.1.86 (low) |
| **GP16** | | P | Acetyl transferase | 0.87 | Iaa acetyltransferase (PDB: 1y9k, Z-score: 13) | *Bacillis subtilis* Acetyltransferase (PDB: 1yvk, E-value:2.73e-6) | n/a | Acid-thiol ligases EC:6.2.1.14 (low) |
| **GP17** | | | | 0.31 | - | - | 1-35 aa IDR, 49-66 aa TM | Carboxy- and carbamoyltransferases<br>EC:2.1.3.1 (low) |
| **GP18** | Structurally similar to PM2 protein p3 | P | Minor capsid protein | T | - | - | | Glycosylases<br>EC:3.2.1.39 (low) |
| **GP19** | Major capsid protein - DJR viral lineage | P | | T | - | - | 2-134 aa, 135-264 aa Viral major capsid protein (IPR056906) | Glycosylases<br>EC:3.2.1.129 (low) |
| **GP20** | | | | 0.42 | - | - | n/a | Oxidoreductases<br>EC:1.12.2.1 (low) |
| **GP21** | Putative ATPase-DNA translocase ftsk | | | 0.93 | NTPase from STIV2 (PDB: 4kfs, Z-score of 19.3) | NTPase from STIV2 (PBD:4kfr, and E-value of 1.74e-12) | 12-90 aa Helicase HerA, central domain;113-188 aa Helicase HerA-like (IPR008571) | Site-specific deoxyribonuclease EC:3.1.21.4 (low) |
| **GP22** | | | | 0.45 | - | - | n/a | Transferase (amino group) EC:5.4.3.3 (low) |
| **GP23** | | P | | 0.55 | - | - | n/a | Decarboxylase<br>EC:7.2.4.3 (low) |

### Supplementary Figures

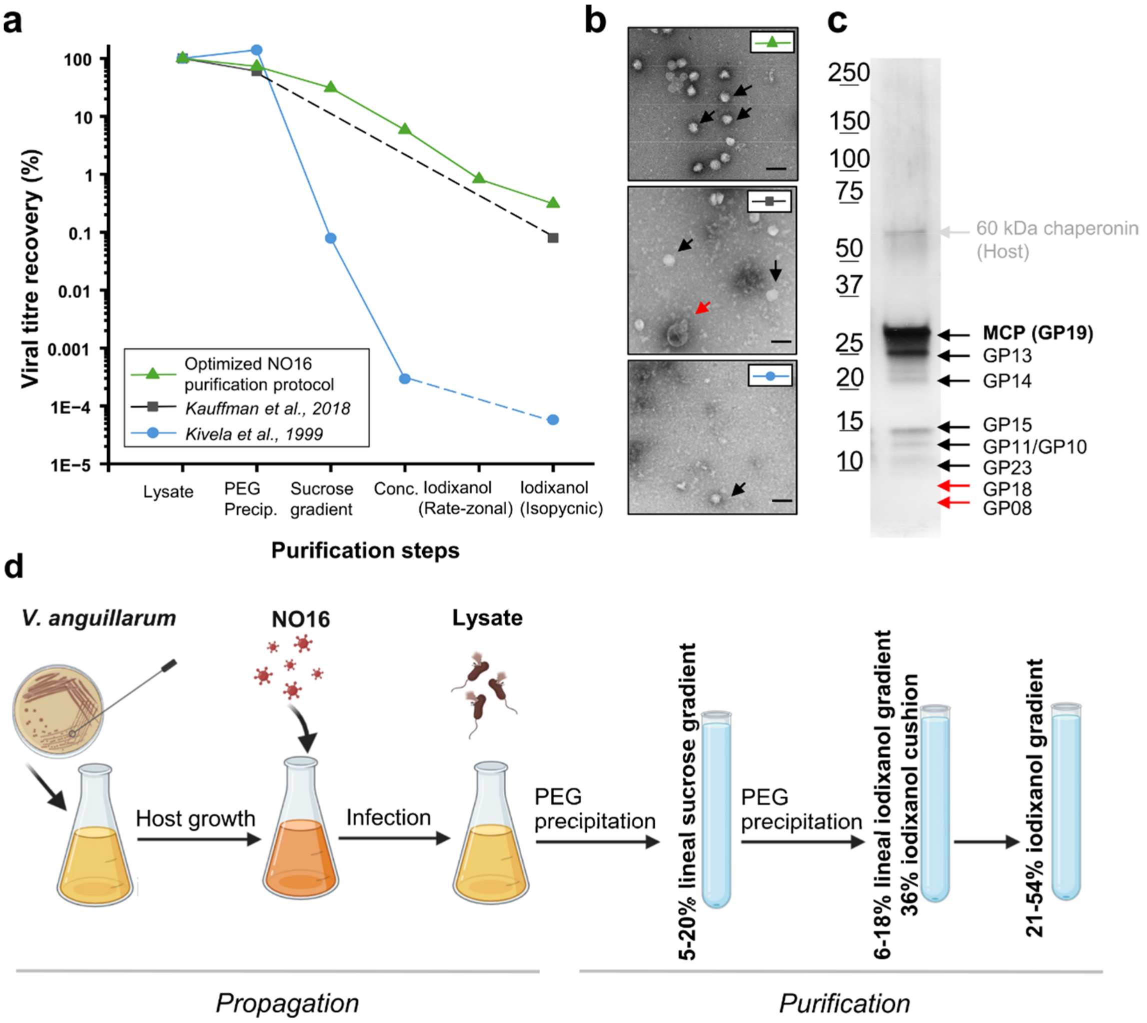

**Figure S1. Optimization of the NO16 purification.** (**a**) Viral titre recovery in each purification step in tested protocols. Dashed lines indicate intermediate steps absent in the protocol. *Conc.* stands for concentration, which was by differential ultracentrifugation in *Kivela et al., 1999* and by PEG precipitation in the optimized protocol. (**b**) Negative staining micrographs of the sample at the last purification step. Black arrows indicate NO16 particles; red arrows indicate host contaminants. Scale bars represent 100 nm. (**c**) SDS-PAGE analysis of NO16 particles purified by the optimized protocol. Tentative identity assignments to each band are based on protein molecular weights. Red arrows indicate the estimated position for non-visible bands (**Table 1, Supplementary File 1b**). The grey arrow indicates the host 60 kDa chaperonin band, confirmed by LC-MS/MS (**Supplementary File 1a**). (**d**) Schematic representation of the propagation and purification workflow for the optimized protocol. Created using Biorender.com.

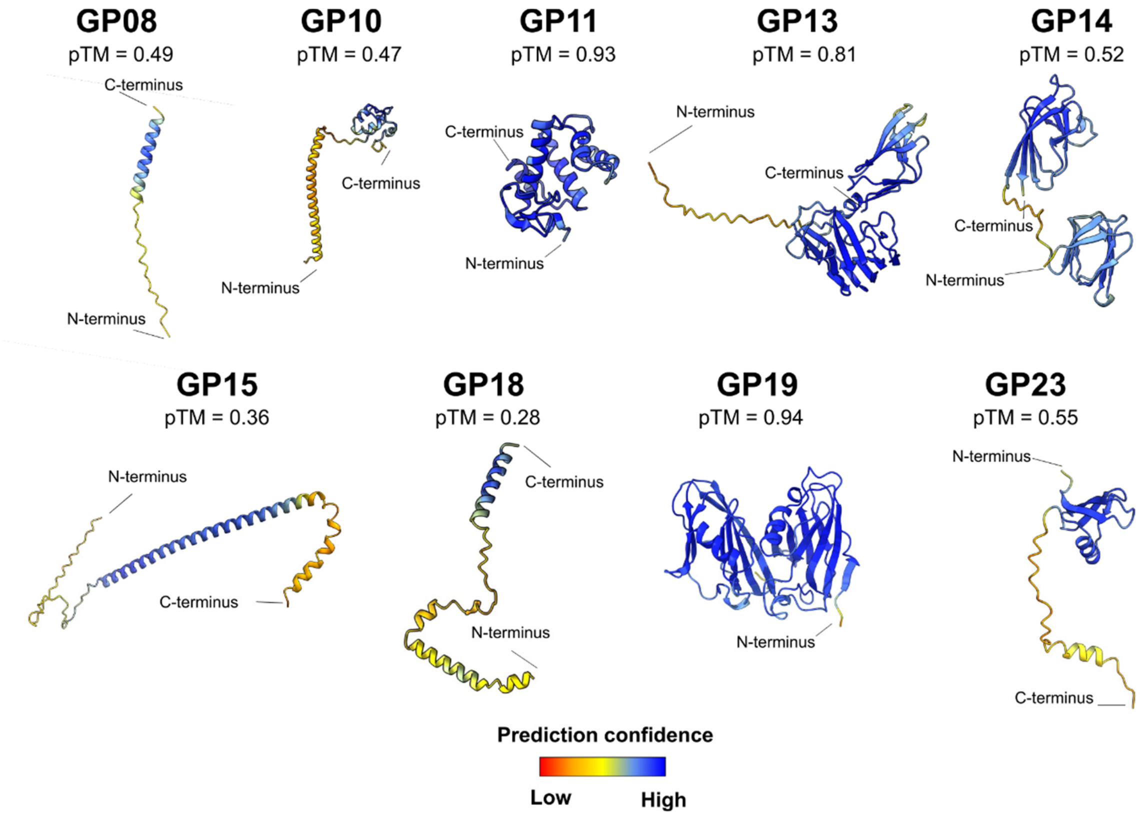

**Figure S2. AlphaFold3 predictions of NO16 proteins detected in purified virions by mass-spectrometry**. Models are coloured by prediction confidence. The pTM score is indicated next to each model.

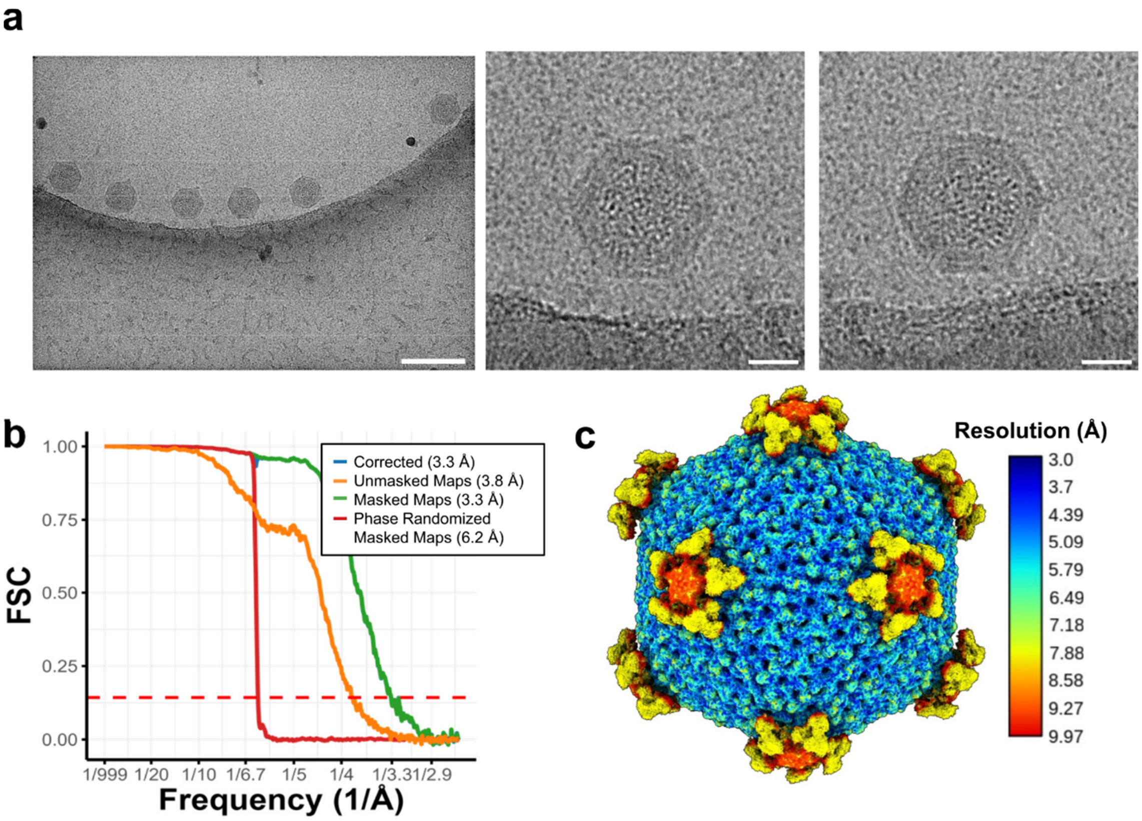

**Figure S3. NO16 cryo-EM data, average and local resolution.** (**a**) Cryo-EM motion corrected micrograph (left), and representative NO16 particles (centre and right). The scale bar represents 100 nm in the micrograph and 20 nm on the particles. (**b**) Fourier Shell Correlation (FSC) curve indicating an average resolution of 3.3 Å in the final map as provided by RELION post-process. Note that the corrected and masked maps curves overlap. (**c**) Cryo-EM map coloured by local resolution, calculated by Xmipp Local MonoRes (Vilas, Gomez-Blanco et al., 2018).

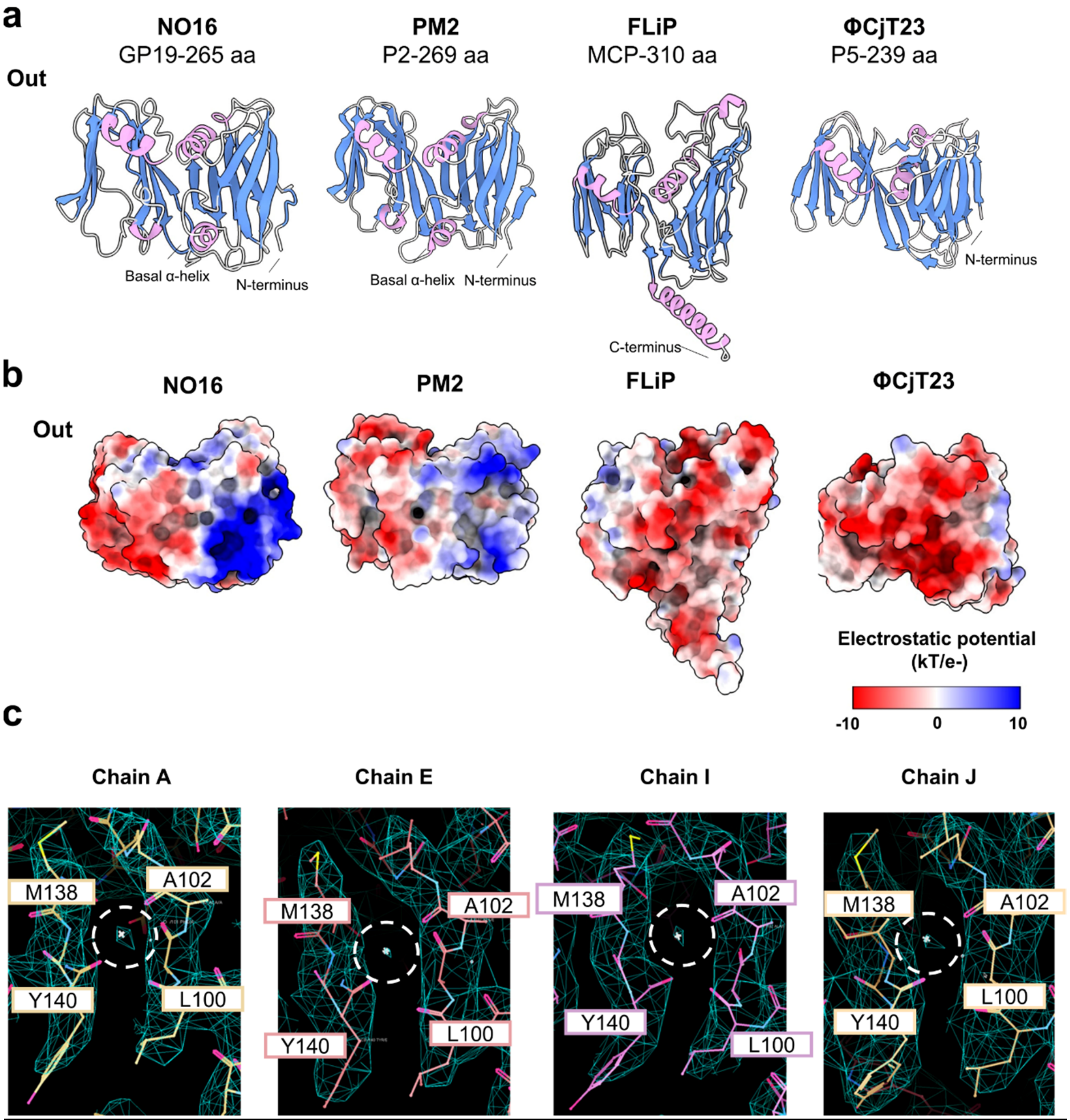

**Figure S4. Comparison of the MCP structure of *p*T = 21 bacteriophages of the DJR lineage.** (**a**) Monomers of NO16, PM2 (PDB: 2vvf), FLiP (PDB: 5oac), and ΦCjT23 (PDB: 7zzz) MCPs with α-helices in pink and β-sheets in blue. Protein names and length in amino acids (aa) are indicated. View from the intra-capsomer side. (**b**) MCP monomers coloured by surface Coulombic electrostatic potential colouring orientated as in (a). (**c**) Cation coordination site at the base of the NO16 MCP visualized in Coot (Emsley, Lohkamp et al., 2010). Map in light blue. Dashed lines circle the cation map density. Residues involved in the interactions are indicated in squares.

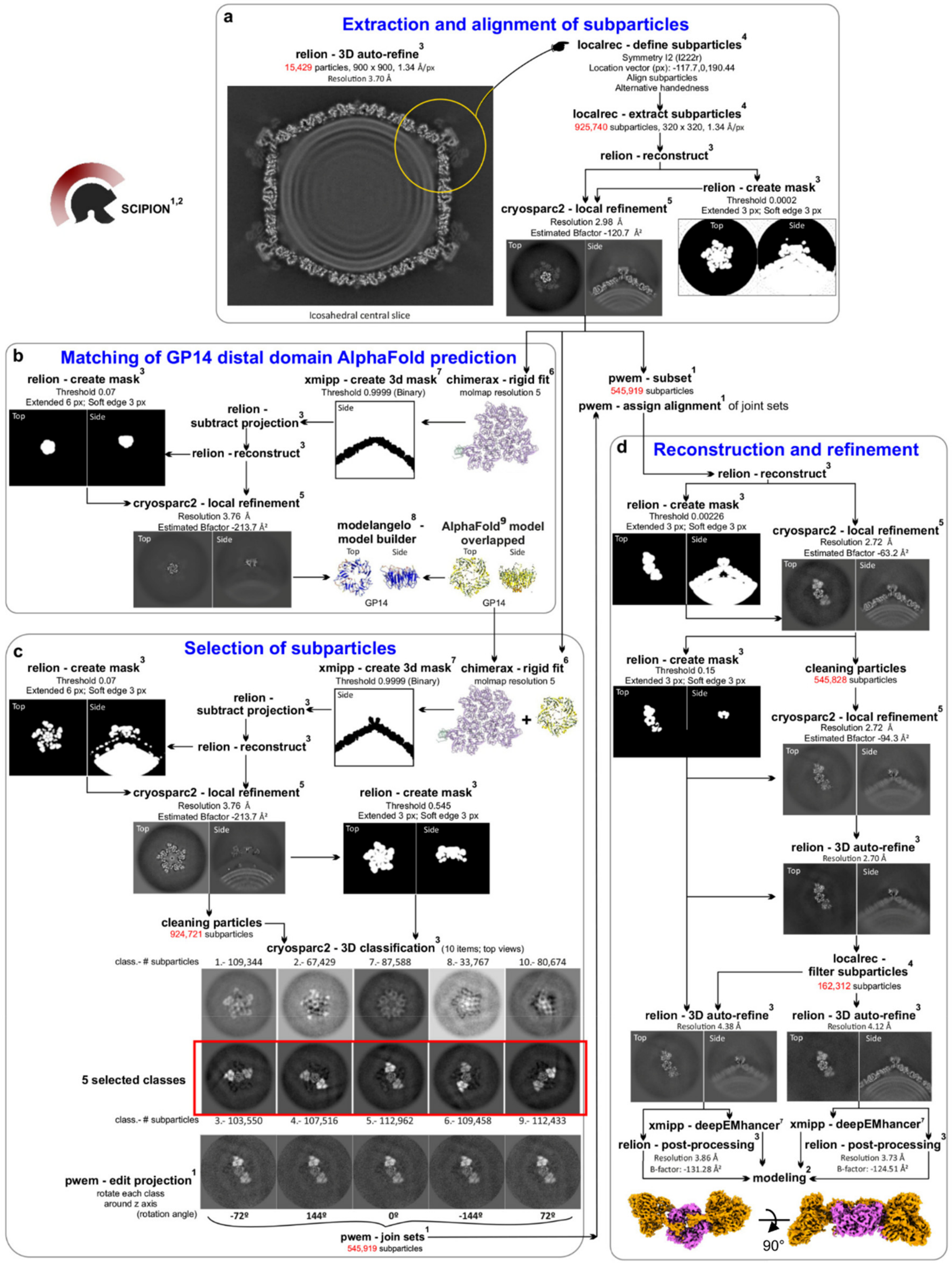

**Figure S5. Local reconstruction workflow of NO16 vertex region.** (**a**) Extraction and alignment of vertex sub-particles from the icosahedral refined map centred at the 5-fold symmetry axis. (**b**) First stage of capsid projection subtraction in the aligned particles to model automatically the distal domain of GP14. This initial GP14 model allows matching the AlphaFold structure prediction to the GP14 distal domain. (**c**) Second stage of capsid projection subtraction in the aligned particles to remove all capsid signal except the five petals surrounding the penton base. Classification without alignment of subtracted particles discriminates the selected oriented pool of particles for further refinement. (**d**) Assignment of subtracted particles orientations to the aligned initial sub-particles to perform the local refinement of the whole vertex (without masking)

and of the distal domain of penton base surrounded by two opposing petals (with local masking). Top image views belong to the slice 180 of the reconstructed map while side views belong to the central slice 160. The entire processing was performed using the Scipion software framework using methods from plugins described in https://scipion.i2pc.es/report_protocols/packages. (1) Scipion (de la Rosa-Trevin, Quintana et al., 2016). (2) Scipion for modelling (Martínez, Jiménez-Moreno et al., 2020). (3) RELION(Scheres, 2012). (4) Localrec (Abrishami, Ilca et al., 2021). (5) CryoSPARC2 (Punjani, Rubinstein et al., 2017). (6) ChimeraX(Meng, Goddard et al., 2023). (7) Xmipp3 (Strelak, Jimenez-Moreno et al., 2021). (8) ModelAngelo(Jamali, Kall et al., 2024). (9) AlphaFold 3 (Abramson, Adler et al., 2024).

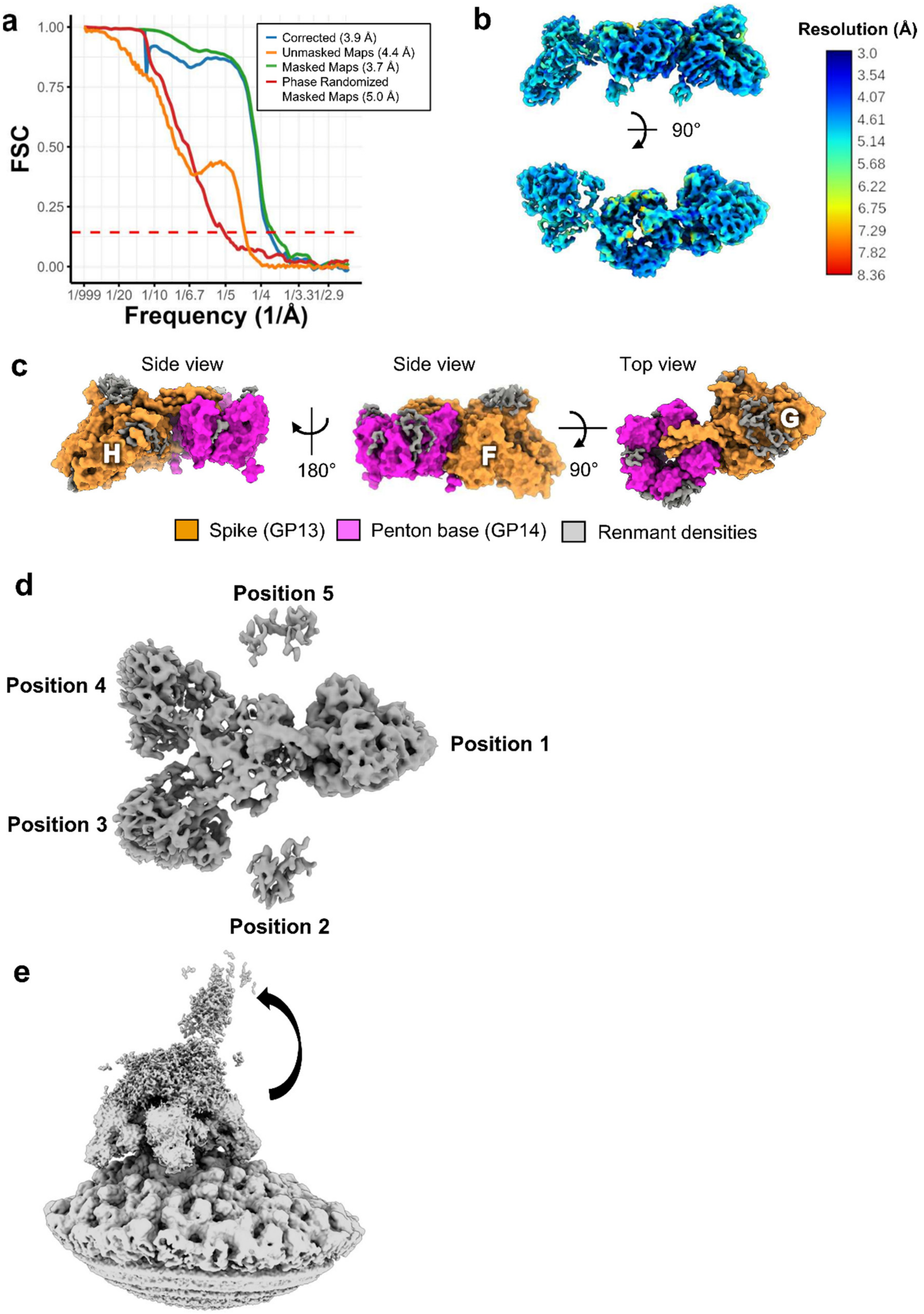

**Figure S6. Local reconstruction of the vertex region.** (**a**) Fourier Shell Correlation (FSC) curve for the NO16 local reconstructed vertex region map, with an average resolution of 3.9 Å, as provided by RELION post-process. (**b**) NO16 local reconstructed cryo-EM map coloured by local resolution, calculated by Xmipp Local Monores. (**c**) Molecular model and remnant densities on the locally reconstructed map, regions in the map where the electron density is present, but the identity of the amino acids could not be assigned unequivocally. (**d**) Local classification map from one spike fixed at position 1, showing density on to opposing spike position 3 and 4. (**e**) Map showing density extending from the penton base cavity region to a nearly perpendicular orientation, probably indicating open conformation of the spike.

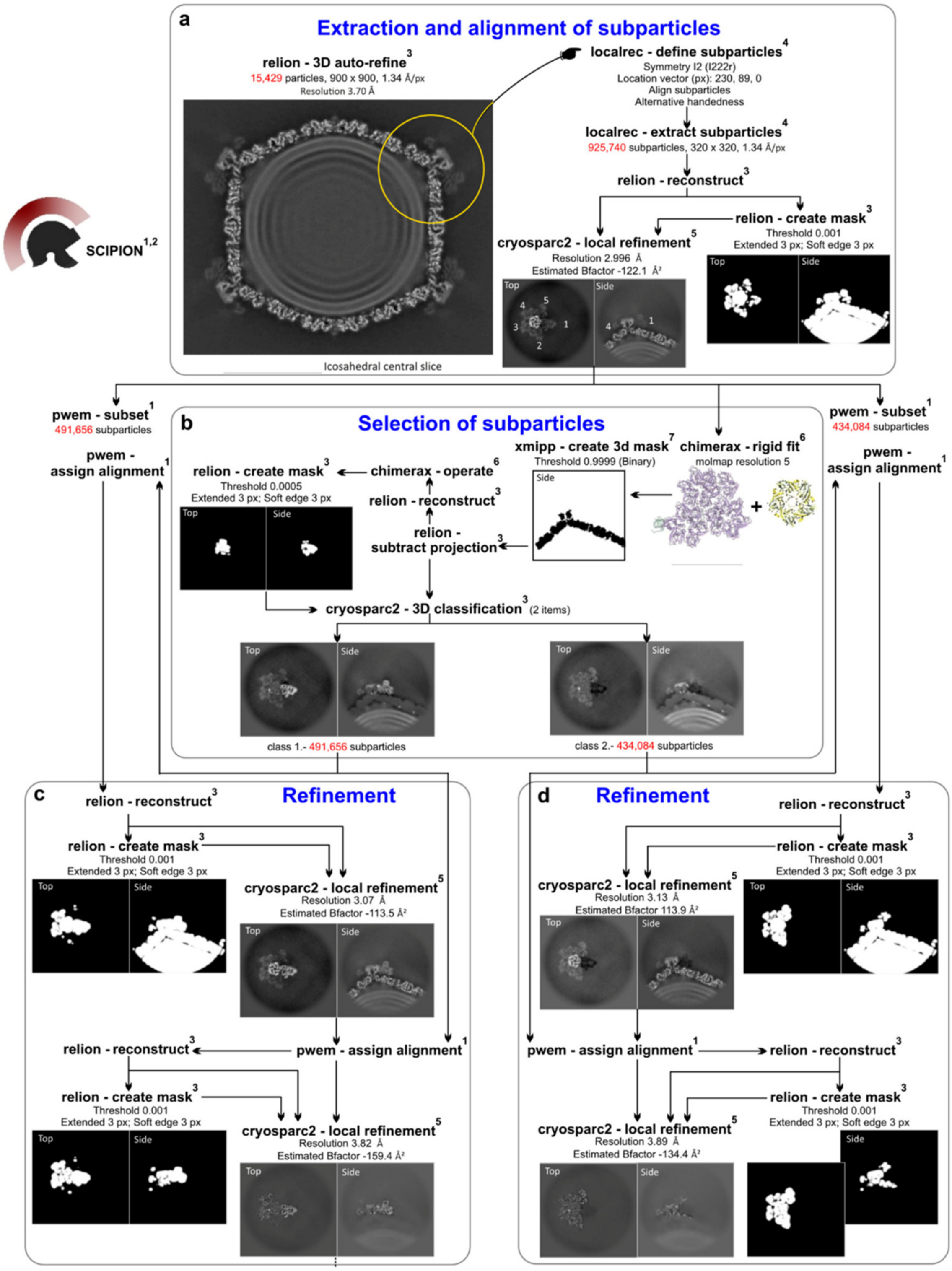

**Figure S7. Workflow of inspecting adjacent positions of a fixed spike in NO16 vertex region.** (**a**) Extraction and alignment of vertex sub-particles from the icosahedral refined map centred in a vertex spike. (**b**) Capsid projection subtraction in the aligned particles to remove any capsid signal except the five petals surrounding the penton base. Classification without alignment of subtracted particles discriminates between presence and absence of spike in position 1. Both sub-particle classes generated are selected for further refinement. (**c, d**) Assignment of subtracted particles' orientations to the aligned initial sub-particles to perform the local refinement of the whole vertex. The refined orientations of sub-particles will be assigned again to subtracted particles just to align locally distal penton base and surrounding spikes. Top image views belong to the slice 154 of the reconstructed map while side views belong to slice 200. The entire processing was performed using the Scipion software framework using methods from plugins described in https://scipion.i2pc.es/report_protocols/packages. (1) Scipion (de la Rosa-Trevin et al., 2016). (2) Scipion for modelling (Martínez et al., 2020). (3) RELION(Scheres, 2012). (4) Localrec (Abrishami et al., 2021). (5) CryoSPARC2 (Punjani et al., 2017). (6) ChimeraX(Meng et al., 2023). (7) Xmipp3 (Strelak et al., 2021).

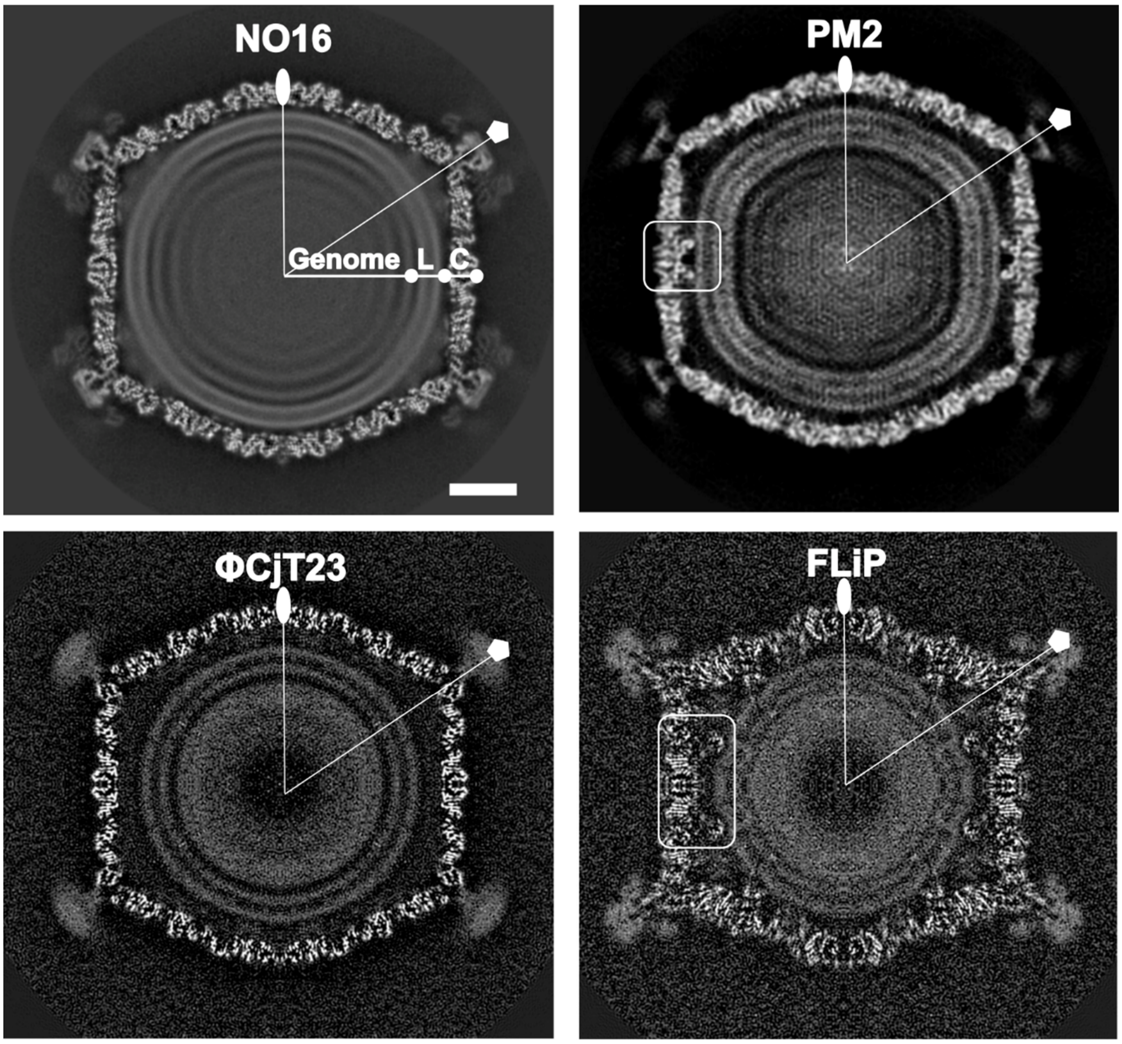

**Figure S8. Map comparison with *p*T = 21 bacteriophages in the lineage.** Central slices of cryo-EM maps for NO16 (this work), PM2 (EMDB:1082), ΦCjT23 (EMDB:15042) and FLiP (EMDB: 3771). C, capsid; L; lipid membrane. The scale bar represents 20 nm. The rounded rectangle indicates capsid-membrane connections.

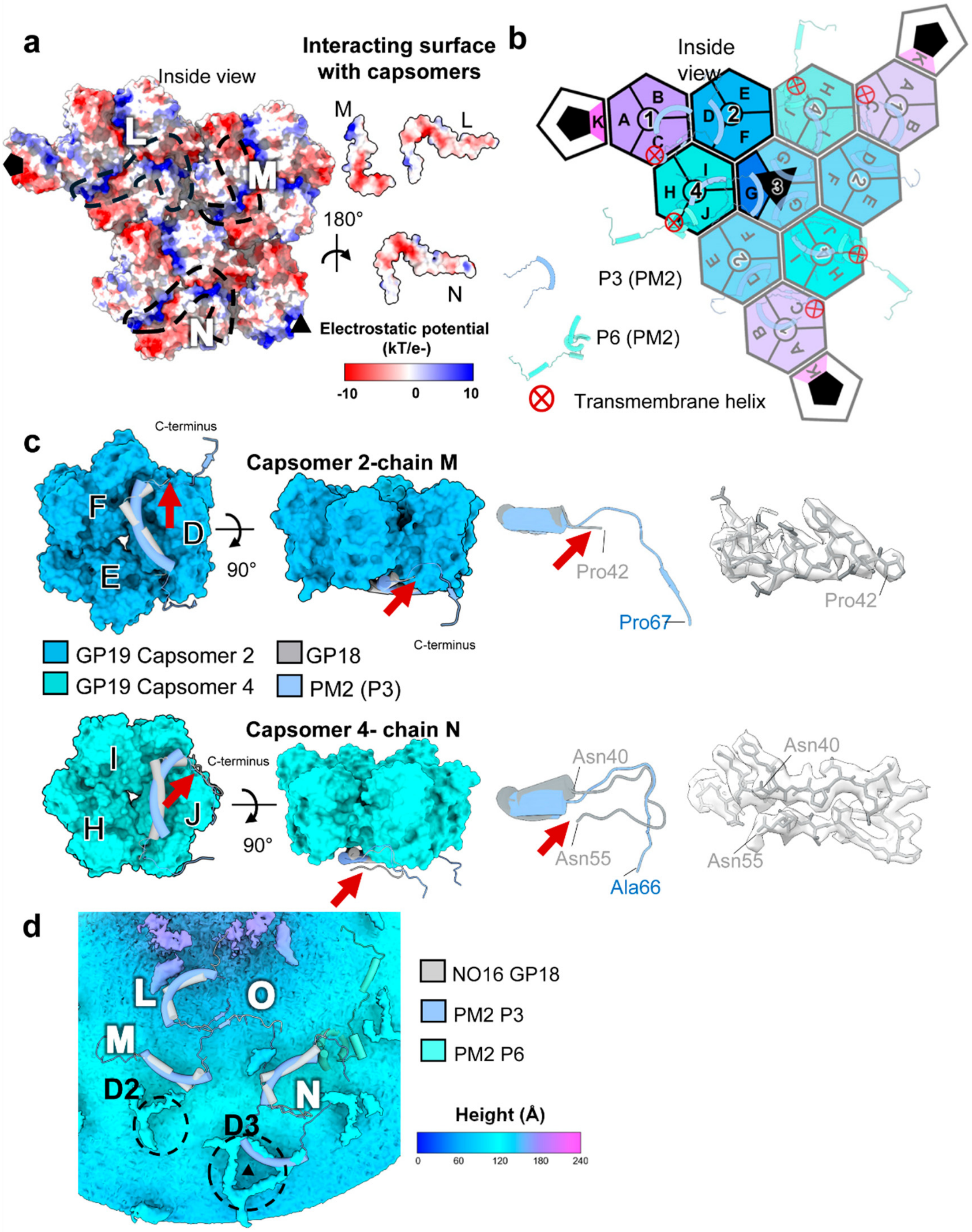

**Figure S9. Minor capsid proteins comparison within NO16 and PM2.** (**a**) Asymmetric unit (AU) from the inside of the capsid coloured by Coulombic electrostatic potential. Left, AU with mCPs removed; their positions are indicated by dashed silhouette outlines. Right, mCPs are coloured by Coulombic electrostatic potential. (**b**) Schematic organization of PM2 P3 and P6 mCPs on top of NO16 facet. (**c**) Capsomers 2 and 4 from inside (left) and side (right), GP18 protein and PM2 P3 overlapped. Red arrows indicate the conformational change on C-terminal of GP18 compared to P3 of PM2. On the right, the molecular model of GP18 chains M and N, along with its corresponding cryo-EM map densities. (**d**) Local reconstructed map showing remnant densities corresponding to mCPs with traced NO16 and PM2 mCPs on top.

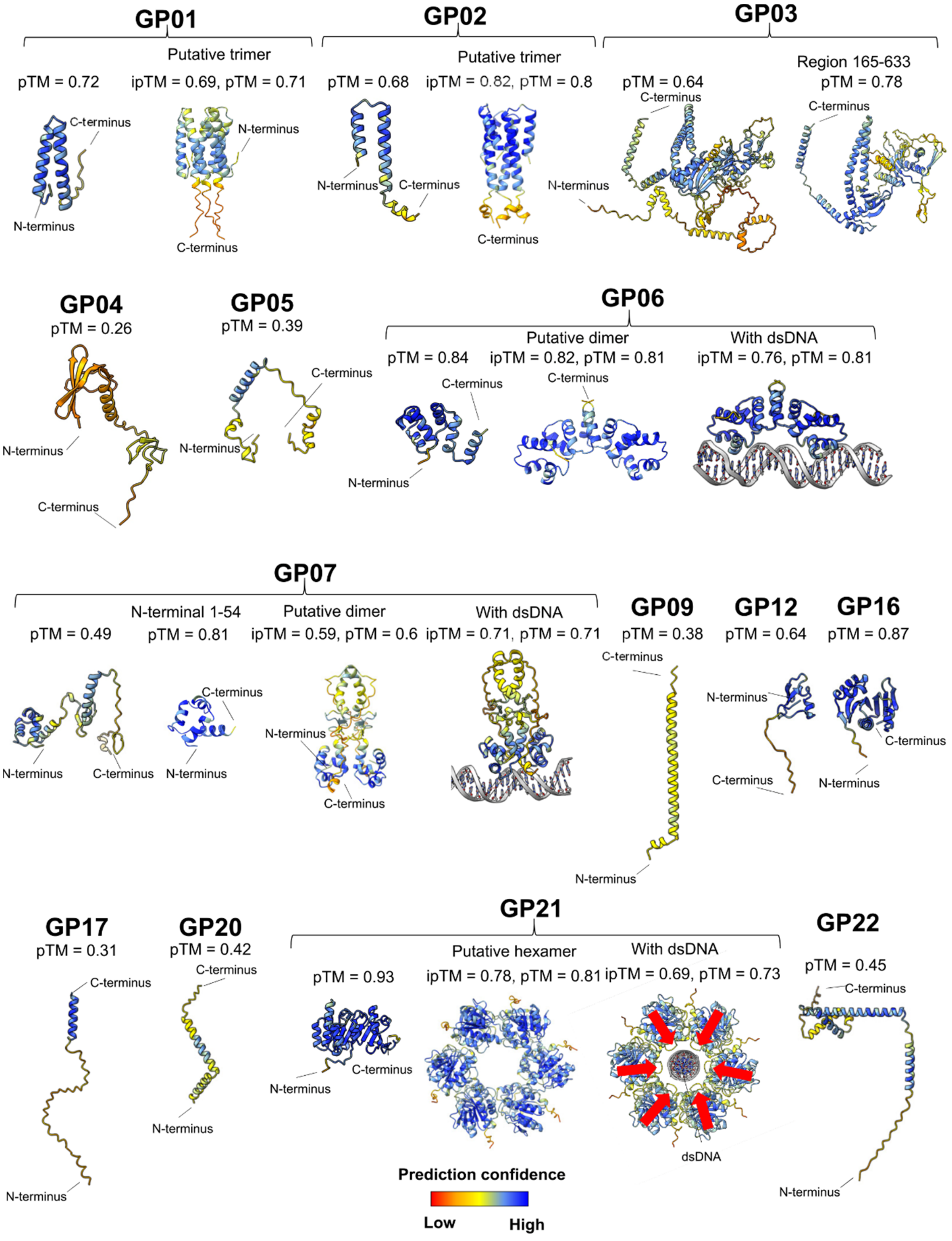

**Figure S10. AlphaFold3 predictions for NO16 encoded proteins not detected in the virion**. See Fig. S2 for proteins detected in the viral particle by mass spectrometry. Models are coloured by prediction confidence. pTM and ipTM values (for oligomers) are indicated next to each model. Red arrows indicate the helicase domain of GP21.

**Supplementary references**

Abramson J, Adler J, Dunger J, Evans R, Green T, Pritzel A, Ronneberger O, Willmore L, Ballard AJ, Bambrick J, Bodenstein SW, Evans DA, Hung CC, O'Neill M, Reiman D, Tunyasuvunakool K, Wu Z, Zemgulyte A, Arvaniti E, Beattie C et al. (2024) Accurate structure prediction of biomolecular interactions with AlphaFold 3. *Nature* 630: 493-500

Abrishami V, Ilca SL, Gomez-Blanco J, Rissanen I, de la Rosa-Trevin JM, Reddy VS, Carazo JM, Huiskonen JT (2021) Localized reconstruction in Scipion expedites the analysis of symmetry mismatches in cryo-EM data. *Prog Biophys Mol Biol* 160: 43-52

de la Rosa-Trevin JM, Quintana A, Del Cano L, Zaldivar A, Foche I, Gutierrez J, Gomez-Blanco J, Burguet-Castell J, Cuenca-Alba J, Abrishami V, Vargas J, Oton J, Sharov G, Vilas JL, Navas J, Conesa P, Kazemi M, Marabini R, Sorzano CO, Carazo JM (2016) Scipion: A software framework toward integration, reproducibility and validation in 3D electron microscopy. *J Struct Biol* 195: 93-9

Emsley P, Lohkamp B, Scott WG, Cowtan K (2010) Features and development of Coot. *Acta Crystallogr D Biol Crystallogr* 66: 486-501

Jamali K, Kall L, Zhang R, Brown A, Kimanius D, Scheres SHW (2024) Automated model building and protein identification in cryo-EM maps. *Nature* 628: 450-457

Krissinel E, Henrick K (2007) Inference of macromolecular assemblies from crystalline state. *J Mol Biol* 372: 774-97

Liebschner D, Afonine PV, Baker ML, Bunkoczi G, Chen VB, Croll TI, Hintze B, Hung LW, Jain S, McCoy AJ, Moriarty NW, Oeffner RD, Poon BK, Prisant MG, Read RJ, Richardson JS, Richardson DC, Sammito MD, Sobolev OV, Stockwell DH et al. (2019) Macromolecular structure determination using X-rays, neutrons and electrons: recent developments in Phenix. *Acta Crystallogr D Struct Biol* 75: 861-877

Martínez M, Jiménez-Moreno A, Maluenda D, Ramírez-Aportela E, Melero R, Cuervo A, Conesa P, del Caño L, Fonseca YC, Sánchez-García R, Strelak D, Conesa JJ, Fernández-Giménez E, de Isidro F, Sorzano COS, Carazo JM, Marabini R (2020) Integration of cryo-EM model building software in Scipion. *Journal of Chemical Information and Modeling* 60: 2533-2540

Meng EC, Goddard TD, Pettersen EF, Couch GS, Pearson ZJ, Morris JH, Ferrin TE (2023) UCSF ChimeraX: Tools for structure building and analysis. *Protein Sci* 32: e4792

Punjani A, Rubinstein JL, Fleet DJ, Brubaker MA (2017) cryoSPARC: algorithms for rapid unsupervised cryo-EM structure determination. *Nat Methods* 14: 290-296

Scheres SH (2012) RELION: implementation of a Bayesian approach to cryo-EM structure determination. *J Struct Biol* 180: 519-30

Strelak D, Jimenez-Moreno A, Vilas JL, Ramirez-Aportela E, Sanchez-Garcia R, Maluenda D, Vargas J, Herreros D, Fernandez-Gimenez E, de Isidro-Gomez FP, Horacek J, Myska D, Horacek

M, Conesa P, Fonseca-Reyna YC, Jimenez J, Martinez M, Harastani M, Jonic S, Filipovic J et al. (2021) Advances in Xmipp for cryo-electron microscopy: from Xmipp to Scipion. *Molecules* 26

Vilas JL, Gomez-Blanco J, Conesa P, Melero R, Miguel de la Rosa-Trevin J, Oton J, Cuenca J, Marabini R, Carazo JM, Vargas J, Sorzano COS (2018) MonoRes: Automatic and Accurate Estimation of Local Resolution for Electron Microscopy Maps. *Structure* 26: 337-344 e4
